# sc-pcQTL: hurdle-based co-expression modeling for multi-gene QTL mapping in single-cell RNA-seq data

**DOI:** 10.64898/2026.08.18.745314

**Authors:** Junkai Zhang, Yi Huang, Melina Claussnitzer, Masahiro Kanai, Wei Zhou

**Author notes:** These authors jointly supervised this work (Masahiro Kanai, Wei Zhou).

## Abstract

**Motivation:** Single-cell expression quantitative trait locus (eQTL) studies can resolve cell-type-specific genetic effects, but conventional gene-by-gene analyses do not directly capture coordinated genetic regulation of neighboring genes. Principal-component QTL (pcQTL) mapping can summarize such multi-gene effects, but existing approaches were developed for bulk expression and are not designed for sparse single-cell counts. Results: We developed sc-pcQTL, a framework that applies two-component hurdle modeling and sliding-window clustering to identify local co-expression clusters, summarizes each cluster using principal components, and maps cis-pcQTLs. In simulations, the individual hurdle components controlled type I error, while the component-union screening rule was substantially more powerful than donor-level pseudobulk correlation tests. Applied to 1.24 million peripheral blood mononuclear cells from 982 OneK1K donors across 10 cell types, sc-pcQTL identified 2,485 local co-expression clusters and conducted QTL mapping for 4,353 cluster-PC phenotypes at single-cell resolution, of which 2,040 had at least one significant cis-pcQTL association. Fine-mapping and colocalization with genome-wide association study loci across 1,163 phenotypes in the FinnGen study identified 394 colocalized QTL-GWAS signal groups. Each group comprised fine-mapped QTL and GWAS signals connected through one or more colocalization links within the same cell type and local gene cluster. Of these groups, 46 were pcQTL-specific and contained no colocalized single-gene eQTL from a constituent gene. Locus-level analyses further revealed cell-type-specific multi-gene regulatory effects. Thus, sc-pcQTL complements conventional single-gene eQTL analysis by identifying trait-relevant regulatory signals shared across neighboring genes. Availability and implementation: The sc-pcQTL software is openly available at https://github.com/ZhouLabGenetics/sc-pcQTL; analysis and figure-generation scripts are available at https://github.com/ZhouLabGenetics/sc-pcQTL_code; and the summary result tables are publicly available on Zenodo (DOI: https://doi.org/10.5281/zenodo.21222687).

**Supplementary material:** Supplementary material accompanies this preprint.

## Introduction

Large-scale transcriptomic studies have identified correlated expression among a substantial fraction of neighboring genes (Ghanbarian and Hurst, 2015; Ribeiro et al., 2021). Several biological mechanisms may underlie this local co-expression among neighboring genes. A shared enhancer can activate multiple distal genes within the same chromatin contact domain (Huang et al., 2025), while genes regulated by the same transcription factor can also colocalize within specialized transcriptional compartments, providing another mechanism for co-expression (Schoenfelder et al., 2010). Single-nucleotide changes can also alter multiple overlapping transcription-factor binding sites and thereby modify regulatory output (Khetan et al., 2025). Consistent with these mechanisms, local co-expression QTLs can affect several nearby genes and are more often associated with multiple human traits than other eQTLs (Ribeiro et al., 2021). These observations motivate modeling local genetic regulation as a multi-gene process rather than as a collection of independent single-gene effects.

Single-cell RNA sequencing (scRNA-seq) provides a cell-resolved setting for studying such multi-gene regulation and can reveal cell-type-specific effects that are masked when expression is averaged across heterogeneous cells in bulk measurements (van der Wijst et al., 2018; Cuomo et al., 2020; Yazar et al., 2022). Standard *cis*-expression quantitative trait locus (*cis*-eQTL) scans test nearby variants against one gene at a time and have been highly successful in identifying regulatory variants (Võsa et al., 2021; Zhou et al., 2024). However, this gene-by-gene framework does not directly model regulatory effects shared across neighboring genes; coordinated local regulatory programs may consequently be missed or fragmented into several weaker signals. Related methods have mapped genotype-dependent changes in pairwise co-expression from scRNA-seq or in broader co-expression networks from bulk-tissue data, but these methods target edges or network structure rather than local multi-gene expression phenotypes (van der Wijst et al., 2018; Li et al.,2023; Kaptijn et al., 2025; Hu et al., 2025).

Recent multi-gene QTL work used principal-component QTL (pcQTL) mapping of co-expressed neighboring genes to identify regulatory associations missed by conventional single-gene analyses. These additional signals also improved colocalization with genome-wide association study (GWAS) loci, supporting pcQTL mapping as an efficient strategy for detecting and interpreting shared local regulatory effects (Lawrence et al., 2026). The existing approach, however, was developed for bulk RNA-seq and is not directly suited to sparse single-cell counts. Variation in sequencing-depth, UMI sampling noise, extreme sparsity, and differences in cell numbers among donors can distort correlation estimates and produce miscalibrated *p*-values when bulk-oriented methods are applied directly (Hafemeister and Satija, 2019; Su et al., 2023). Pseudobulk analysis^1^ provides donor-level summaries resembling bulk RNA-seq data, but averaging can attenuate cell-state-specific expression structure and reduce power to detect regulatory effects confined to particular cellular states (Zimmerman et al.,2021). A framework is therefore needed that connects sparse single-cell co-expression testing with local module construction and downstream multi-gene *cis*-QTL mapping.

To address this gap, we developed sc-pcQTL, a hurdle-based framework that links single-cell co-expression testing, local multi-gene feature construction, and *cis* principal-component QTL (*cis*-pcQTL) mapping (Mullahy, 1986).

## Results

### Overview of sc-pcQTL

sc-pcQTL comprises four stages, each applied separately within a cell type (Figure 1). First, a two-component hurdle model tests pairwise co-expression among eligible gene pairs on the same chromosome. The detection component tests association in zero-versus-nonzero expression, whereas the count component tests association in expression abundance among cells with positive counts. Second, a sliding-window procedure groups nearby genes connected by significant pairwise associations into local co-expression clusters. Third, principal-component analysis summarizes the joint expression of each cluster as one or more cluster-PC phenotypes. Finally, SAIGE-QTL (Zhou et al., 2024) maps *cis* associations between these phenotypes and donor genotypes to identify *cis*-pcQTL associations. We evaluated the empirical type I error and power of the pairwise hurdle screen using two complementary simulation designs and then applied the complete workflow to 1.24 million OneK1K peripheral blood mononuclear cells (PBMCs) from 982 donors across 10 cell types with at least 10,000 cells each (Yazar et al., 2022). We then fine-mapped the identified pcQTLs and matched single-gene eQTLs, tested both QTL types for colocalization with GWAS loci of 1,163 phenotypes in the FinnGen study (Kurki et al., 2023), and quantified enrichment for disease heritability. Together, these analyses assessed whether cluster-PC phenotypes captured trait-associated regulatory signals beyond those represented by matched single-gene eQTLs.

**Figure 1.**
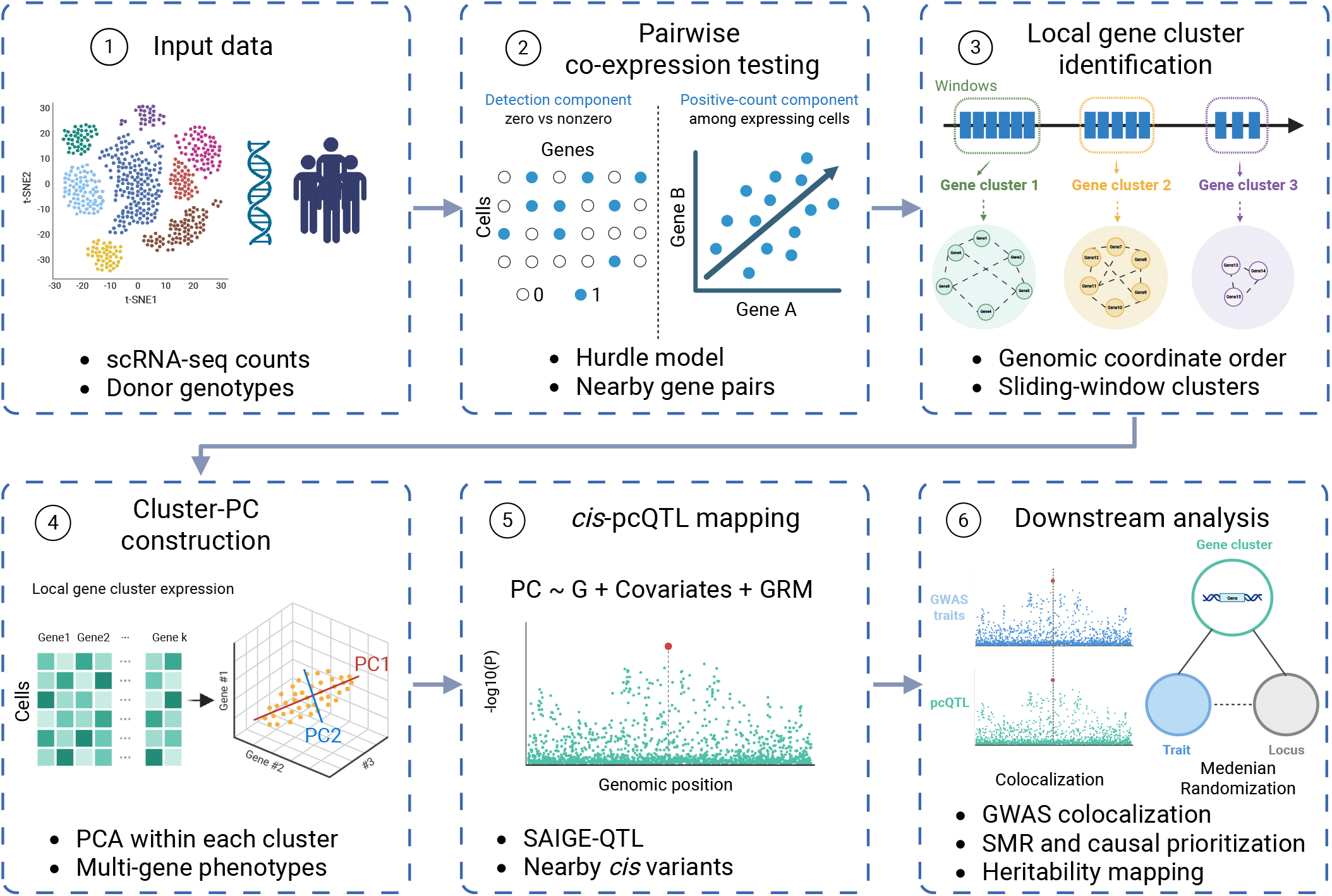
Overview of sc-pcQTL. (1) Single-cell RNA-seq counts and donor genotypes are assembled within each cell type. (2) Pairwise hurdle models test co-expression through the detection and count components. (3) Significant pairwise associations and genomic order define local gene clusters. (4) Principal-component analysis converts cluster expression into multi-gene PC phenotypes. (5) SAIGE-QTL tests nearby variants for *cis*-pcQTL associations. (6) Downstream analyses include GWAS colocalization, summary-data-based Mendelian randomization, and heritability enrichment. Figure created under Agreement No. MI29ZCX6PV.

### Simulation-based evaluation of sc-pcQTL

#### The sc-pcQTL hurdle model was well calibrated and improved power in model-based simulations

We evaluated pairwise co-expression tests using model-based simulations that generated gene detection and positive counts separately (Methods, Section 3.3.1). The hurdle model tested the two expression components individually. For comparison, pseudobulk Pearson and Spearman tests assessed correlations between donor-level mean expression values. We also evaluated a component-union rule that classified an ordered gene-pair test as positive if either component *p*-value was below the nominal threshold.

Under the null, both hurdle components had well-controlled empirical type I error rates (Figure 2a). At *α* = 0.01, the type I error rates for the count and detection components were 0.00962 and 0.00983, respectively. In contrast, the pseudobulk Pearson correlation test was anti-conservative, with a type I error rate of 0.0914, and the Spearman correlation test showed mild inflation, with a type I error rate of 0.0169.

**Figure 2.**
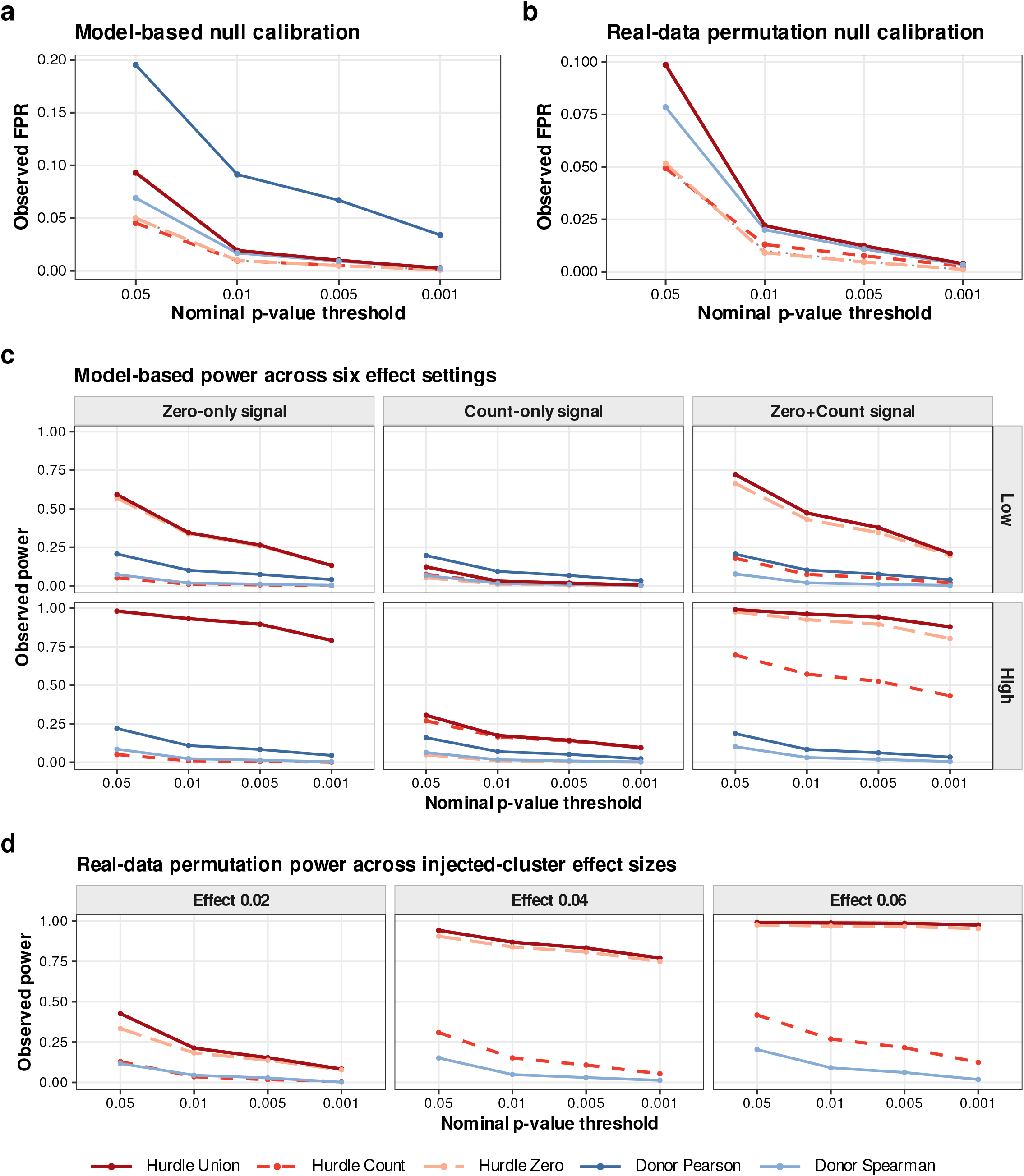
Pairwise hurdle co-expression calibration and power. (a) Model-based simulation null false-positive rates across nominal thresholds. (b) Real-data permutation-based simulation null false-positive rates using a gene-wise-permuted OneK1K count matrix. (c) Model-based power across six settings: detection-only (shown as “Zero-only”), count-only, and joint detection-and-count (shown as “Zero+Count”), each at low and high strength. “Hurdle Zero” denotes the binomial detection component. (d) Real-data permutation-based simulation power across injected cluster-effect coefficients. Panels (a,c) show scenario means across model-based simulation replicates. Panels (b,d) show empirical proportions across eligible tests in the fixed permuted matrix and datasets with injected effects. The component-union rule calls an ordered test positive when either hurdle component passes. Pseudobulk Pearson and Spearman correlations were computed from donor-level mean expression.

In power simulations, the sc-pcQTL hurdle screen substantially outperformed the pseudobulk correlation tests. The largest gains occurred when the shared simulated signal affected only the probability of gene detection (Figure 2c). At *α* = 0.01, power for the detection component reached 0.931 in the high detection-only setting, compared with 0.108 for pseudobulk Pearson and 0.0229 for pseudobulk Spearman. In the high joint setting, in which the same latent signal affected both detection and positive-count levels, the component-union rule achieved 0.962 power, whereas the Pearson and Spearman tests reached 0.0833 and 0.0306, respectively.

#### sc-pcQTL remained well calibrated and more powerful than the baseline method in real-data permutation-based simulations

To evaluate performance under realistic single-cell count distributions, we designed a permutation-based simulation that preserved the sparsity and marginal count distributions observed in OneK1K (Methods, Section 3.3.2). After independently permuting each gene across cells, the empirical type I error rates at *α* = 0.01 were 0.0130 for the count component and 0.00915 for the detection component (Figure 2b). As expected for the unadjusted union of two component tests, the component-union false-positive rate was 0.0220, whereas the corresponding rate for pseudobulk Spearman was 0.0201. After shared cluster effects were introduced at log-scale coefficients of 0.02, 0.04, and 0.06, component-union power was 0.213, 0.869, and 0.988, respectively, compared with 0.0446, 0.0487, and 0.0908 for pseudobulk Spearman (Figure 2d). Thus, the individual hurdle components remained close to their nominal type I error rates, while the component-union achieved substantially greater power than the pseudobulk Spearman method.

An additional simulation evaluated whether introducing within-donor dependence affected null calibration. Rejection rates did not consistently increase relative to matched simulations without donor dependence at the evaluated significance levels (Supplementary Figure S3).

### Application of sc-pcQTL to the OneK1K cohort

#### sc-pcQTL identified local co-expression clusters with shared functional and regulatory features

Applying sc-pcQTL to the 10 primary OneK1K cell types identified 2,485 local co-expression clusters, comprising 5,313 gene occurrences across cell types (Figure 3a,b). CD4^+^ naïve and central-memory T cells (CD4 NC) had the largest number of clusters (513), followed by CD8^+^ naïve and central-memory T cells (CD8 NC; 360), natural killer cells (NK; 340), and CD8^+^ T cells with an effector-memory phenotype (CD8 ET; 316). Nonclassical monocytes (Mono NC) had the fewest clusters among the eligible cell types (101). The median cluster size was two genes, indicating that most clusters represented local gene pairs. Differences in cluster counts across cell types likely reflected variation in both cell number and the number of genes passing the 1% nonzero-expression threshold.

Compared with size-matched sets of neighboring genes, the detected clusters were enriched for shared functional and local regulatory annotations and were less likely to span CTCF peaks or topologically associating domain boundaries (Supplementary Figure S6).

**Figure 3.**
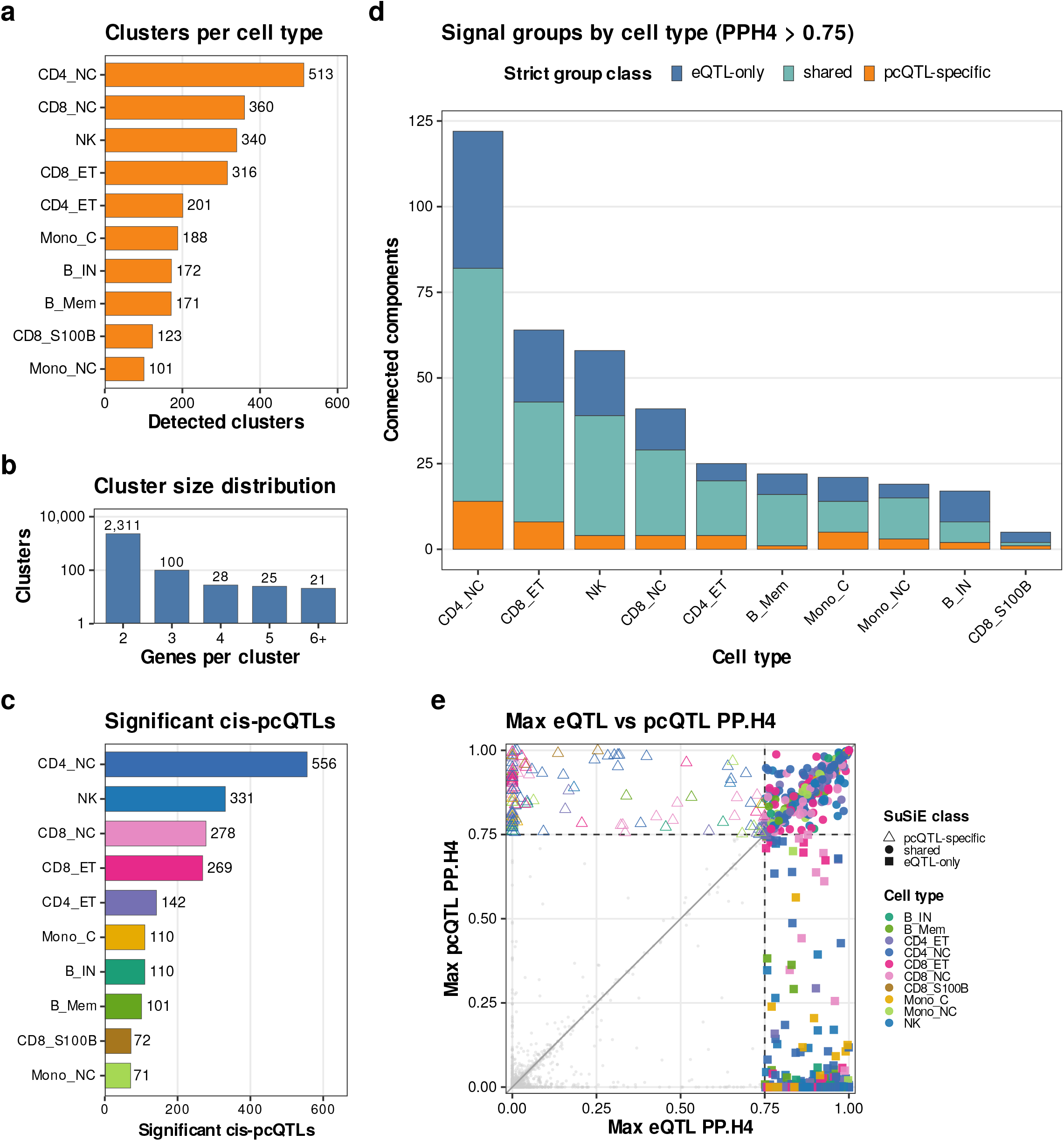
sc-pcQTL cluster landscape, *cis*-pcQTL mapping, and fine-mapped colocalization results in the 10 primary cell types. (a) Number of local co-expression clusters per cell type across 2,485 clusters and 5,313 gene-cell-type assignments. (b) Distribution of cluster sizes pooled across cell types; the count axis is log-scaled, and clusters are predominantly gene pairs. (c) Number of cluster-PC phenotypes with at least one significant *cis*-pcQTL association after within-phenotype Benjamini-Hochberg correction (*q <* 0.05), by cell type. (d) Classes of signal groups by cell type at the primary threshold (PPH4 > 0.75). (e) Maximum single-gene eQTL-GWAS versus pcQTL-GWAS PPH4 by cluster-trait group. Dashed lines mark PPH4 = 0.75; colors denote cell types, shapes denote signal-group classes, and grey points denote non-colocalized groups. Canonical OneK1K labels correspond to the following cell types: immature and naïve B cells (B IN); memory B cells (B Mem); CD4^+^ effector-memory/TEMRA cells (CD4 ET); CD4^+^ naïve and central-memory T cells (CD4 NC); CD8^+^ T cells with an effector-memory phenotype (CD8 ET); CD8^+^ naïve and central-memory T cells (CD8 NC); CD8^+^ T cells expressing *S100B* (CD8 S100B); classical monocytes (Mono C); nonclassical monocytes (Mono NC); and natural killer cells (NK).

Because cross-mappable reads can generate spurious co-expression and eQTL signals (Saha and Battle, 2018; Warmerdam et al., 2026), we compared pcQTL discovery between clusters with and without cross-mappable gene pairs. The proportion of cluster-PC phenotypes with at least one fine-mapped QTL signal did not differ significantly between the groups. Moreover, after excluding cross-mappable clusters, 87.2% of significant pcQTL phenotypes were retained, and most downstream colocalization classifications remained unchanged. We therefore retained these clusters in the primary analysis (Supplementary Figure S7).

#### sc-pcQTL identified cis-pcQTL association signals across 10 OneK1K cell types

We performed principal component analysis (PCA) (Pearson, 1901) on the expression counts of genes in each local co-expression cluster and tested the resulting cluster-derived PC for *cis*-pcQTL associations (Methods, Section 3.5). Of 4,353 successfully tested phenotypes, 2,040 (46.9%) had at least one significant association (within-phenotype FDR *<* 0.05; Figure 3c). The number of significant phenotypes ranged from 71 in Mono NC cells to 556 in CD4 NC cells. NK cells, CD8 NC cells, and CD8 ET cells contributed 331, 278, and 269, respectively.

#### sc-pcQTLs identified additional QTLs colocalized with GWAS beyond single-gene eQTLs

To assess whether cluster-PC phenotypes captured trait-associated regulatory signals beyond those detected by single-gene analysis, we used the Sum of Single Effects (SuSiE) model (Wang et al., 2020) to fine-map *cis*-pcQTLs and the *cis*-eQTLs of constituent genes. We then evaluated colocalization between the fine-mapped QTL signals and fine-mapped GWAS signals in the FinnGen study using `coloc.susie` (Wallace, 2021). We defined colocalized QTL-GWAS signal groups (hereafter referred to as signal groups) as sets of fine-mapped QTL and GWAS signals connected through one or more pairwise colocalization links within the same cell type and local gene cluster. Each retained signal group contained at least one GWAS signal and was classified as eQTL-only, shared, or pcQTL-specific according to its QTL composition (Methods, Section 3.7).

At the primary PPH4 > 0.75 threshold, we identified 394 signal groups: 126 eQTL-only, 222 shared, and 46 pcQTL-specific. The 46 pcQTL-specific signal groups represented a 13.2% increase over the 348 signal groups containing a colocalized single-gene eQTL signal (Figure 3d).

Most contributing cluster-PC phenotypes were dominated by one constituent gene in their PC loadings (Supplementary Table S5). However, their PIP-weighted nominal effects extended to multiple constituent genes rather than being confined to the top-loading gene (Supplementary Figure S10).

Sensitivity analyses using alternative PPH4 thresholds and the single-causal-variant `coloc.abf` model (Giambartolomei et al., 2014) yielded the same overall pattern of eQTL-only, shared, and pcQTL-specific signal groups (Supplementary Figures S8 and S9).

At the cluster-trait level, the maximum eQTL-GWAS and pcQTL-GWAS PPH4 values recapitulated the three signal-group classes: eQTL-only combinations showed stronger evidence for single-gene eQTL colocalization, pcQTL-specific combinations showed stronger evidence for pcQTL colocalization, and shared combinations showed evidence for both QTL types (Figure 3e).

Stratified LD score regression (S-LDSC) (Bulik-Sullivan et al., 2015; Finucane et al.,2015; Gazal et al., 2017) showed that pcQTL-associated variants were enriched for disease heritability. After adjustment for baseline-LD annotations, the pcQTL coefficient remained positive in the separate model but did not reach statistical significance; it was further reduced in the joint pcQTL-eQTL model (Supplementary Methods; Supplementary Figure S11; Supplementary Table S6).

#### Representative pcQTL loci illustrated links between multi-gene regulation and complex traits

The *GIMAP* locus in CD8 NC cells illustrates a biologically informative and representative example of a pcQTL-specific signal. The *GIMAP* PC3 pcQTL in CD8 NC cells colocalized with a lymphocyte-count GWAS locus in the FinnGen study(Kurki et al., 2023), with strong support for a shared causal signal (PPH4 = 0.933) and limited support for distinct causal signals (PPH3 = 0.067; Figure 4). No single-gene eQTL for a constituent gene exceeded the primary colocalization threshold.

**Figure 4.**
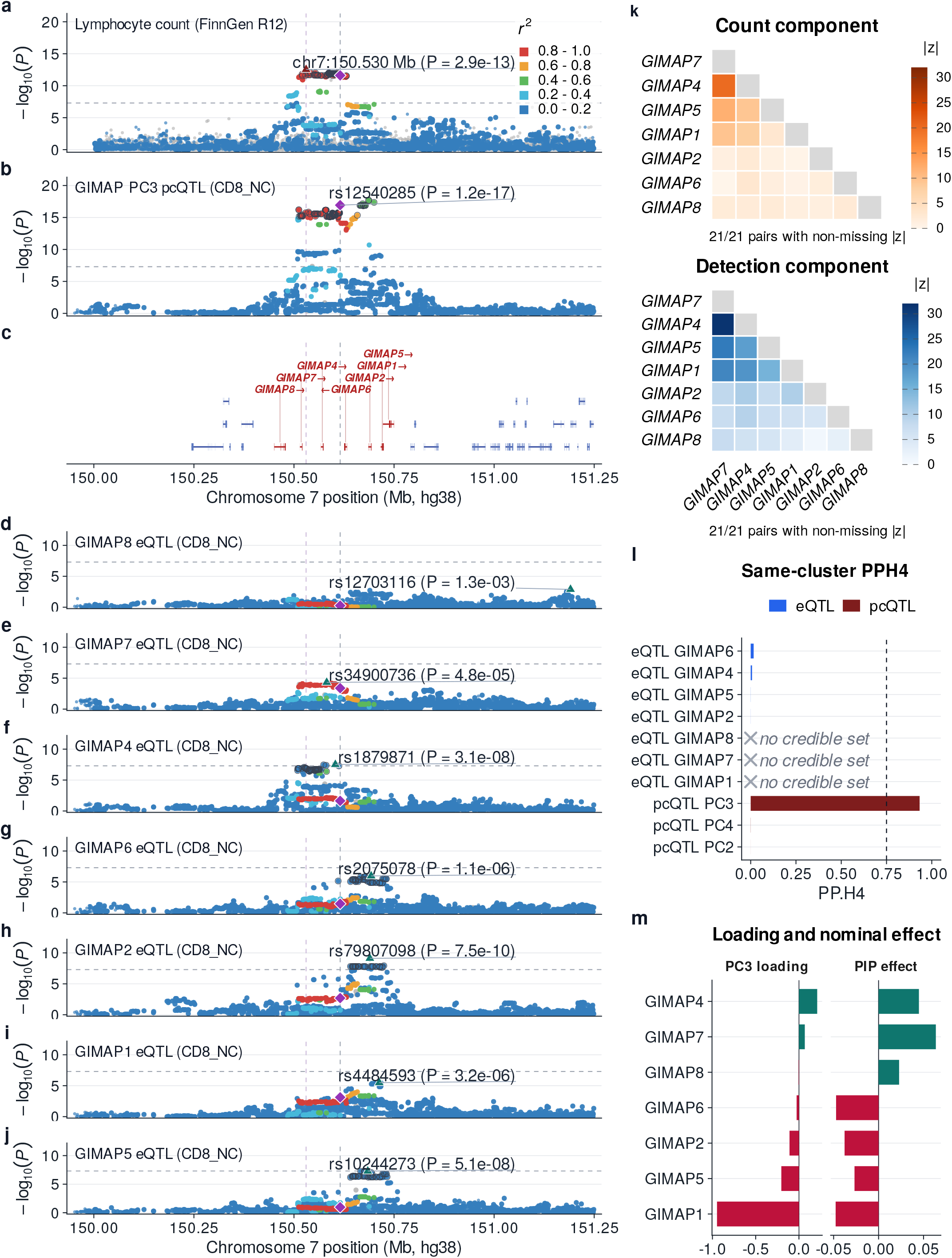
Regional colocalization of the *GIMAP* PC3 pcQTL with lymphocyte count. The pcQTL in CD8^+^ naïve and central-memory T cells (CD8 NC) colocalized with the lymphocyte-count GWAS signal in the FinnGen study(PPH4 = 0.933). Panel descriptions are provided with the continued legend on the following page. (a) Regional association for the lymphocyte-count GWAS in the FinnGen study. (b) Regional association for the OneK1K *GIMAP* PC3 pcQTL in CD8 NC cells. (c) hg38 gene track; *GIMAP* cluster genes are red, other genes are blue, and arrows indicate transcriptional direction. (d–j) Regional single-gene eQTL associations for the seven constituent *GIMAP* genes in the same cell type. (k) Absolute *z*-statistics from the count and detection components of the hurdle model for all 21 within-cluster gene pairs; grey diagonal tiles denote self-comparisons. (l) Posterior probability of a shared causal variant (PPH4) between each same-cluster eQTL or pcQTL signal and GWAS signal in the FinnGen study. The dashed line marks the primary PPH4 = 0.75 threshold; grey crosses identify QTL phenotypes without a SuSiE credible set, for which colocalization was not tested. (m) PC3 gene loadings and PIP-weighted nominal gene effects. Panels a and b share one *™* log_10_ *P* scale, whereas panels d–j share a second scale and have uniform dimensions. All regional association panels use the same hg38 coordinate range and identify the panel-specific lead variant and its *P*-value. Point colors show LD *r*^2^ with the pcQTL lead in the 1000 Genomes Phase 3 European display reference; grey points lack display-LD estimates. Purple diamonds mark the LD reference variant, upward triangles mark panel-specific lead variants, and open dark rings mark variants in SuSiE 95% credible sets.

Pairwise hurdle model results supported co-expression for 16 of the 21 gene pairs (Figure 4). The *GIMAP* cluster did not meet the cross-mappability criterion: all seven genes mapped uniquely to GENCODE v19, and none of the 21 evaluable pairs appeared in the resource’s nonzero-pair table (Supplementary Table S3). Targeted summary-data-based Mendelian randomization (SMR) analysis (Zhu et al., 2016) provided gene-level follow-up within the module, with *GIMAP7* showing the strongest HEIDI-consistent association with lymphocyte count (Supplementary Figure S12d,e). Together, these findings were consistent with distributed multi-gene regulation.

The seven genes in the cluster are neighboring members of the immune-associated *GIMAP* small-GTPase family at 7q36.1. Experimental studies have implicated *GIMAP5* in peripheral T-cell survival and in natural killer- and natural killer T-cell development, *GIMAP1* in mature B- and T-cell development, and *GIMAP6* in T-cell maintenance and autophagy (Limoges et al., 2021; Schulteis et al., 2008; Saunders et al., 2010; Pascall et al., 2018). These established immune functions provide biological context for the association between the *GIMAP* module, the T-cell-specific regulatory signal, and lymphocyte count.

Across the 10 primary cell types, the *GIMAP* cluster-PC pcQTL exceeded the PPH4 > 0.75 threshold only in CD8 NC cells; the next-highest value was PPH4 = 0.66 in CD4 ET cells (Supplementary Figure S12a). Although the complete seven-gene module was also identified as a single local cluster in CD8 ET cells, CD4 NC cells, and NK cells (Supplementary Figure S12b), only the cluster-PC phenotype in CD8 NC cells met the primary colocalization threshold. Thus, recovery of the same co-expression module across cell types did not imply that its cluster-PC phenotype would colocalize with lymphocyte count in every cell type.

A donor-level pseudobulk mixture simulation further showed that the trait-colocalized multi-gene signal was attenuated under the observed cell-type composition and strengthened when CD8 NC cells were isolated or upweighted (Supplementary Figure S12c).

Additional pcQTL-specific examples for the *ASCL2*-*C11orf21*-*TSPAN32, EIF3K*-*ACTN4*, and *EVI2B*-*EVI2A* clusters are shown in Supplementary Figures S13–S15.

## Methods

### Data preprocessing

We analyzed cell-type-specific single-cell expression counts, donor genotypes, and covariates from 982 OneK1K donors (Yazar et al., 2022). To reduce instability in underpowered cell-level co-expression analyses, we excluded cell types with fewer than 10,000 cells before pairwise screening. Ten of the 14 annotated cell types met this threshold, comprising 1,241,711 cells. Expression analyses used OneK1K counts processed with SCTransform, a normalization and variance-stabilization method for single-cell UMI data (Hafemeister and Satija, 2019). Within each eligible cell type, genes expressed in fewer than 1% of cells were excluded from pairwise testing and the same filtered gene set was used for pseudobulk pair screening.

### Hurdle model

Single-cell gene-gene associations were estimated using the two-part hurdle model implemented in *fasthurdle* (Mullahy, 1986; Kanai et al., 2025). The model combined regression of positive expression counts (the count component) with binomial regression of nonzero expression (the detection component). A two-part hurdle framework was previously introduced for testing associations between chromatin-accessibility peaks and gene expression in single-cell multiome data(Liu et al., 2025). We used Poisson rather than Negative-Binomial regression for the count component because the Poisson model was better calibrated for the SCTransform-corrected OneK1K counts (Supplementary Figure S4). The detection component is labeled the zero component in *fasthurdle* outputs and simulation files; we use detection component throughout.

For response gene *Y* and predictor gene *X*, let *D* = **1**(*Y* > 0), let ***Z*** denote the covariate matrix, and let *L* denote log library size. The count component was

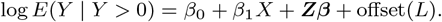

The detection component was

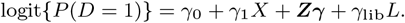

The count component tested the null hypothesis *β*_1_ = 0 for association with positive expression abundance. The detection component tested *γ*_1_ = 0 for association with whether the response gene was detected (*Y* > 0). Both tests were two-sided.

In the OneK1K analysis, ***Z*** included age, sex, genotype principal components 1–6, and PEER factors 1–2, and *L* was the log total read count. The genotype principal components accounted for donor-level ancestry structure (Price et al., 2006), and the PEER factors captured latent expression variation (Stegle et al., 2012).

The pairwise hurdle models treated cells as observations and adjusted for donor-level covariates and library size. For each unordered pair {*g*_1_, *g*_2_}, we fitted both ordered response-predictor directions, *g*_1_ | *g*_2_ and *g*_2_ | *g*_1_, and retained the smaller directional *p*-value separately for the count and detection components. A pair entered cluster construction if either component passed the Bonferroni threshold 0.05*/M*, where 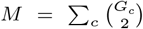 is the total number of tested unordered gene pairs within each autosome and *G*_*c*_ is the number of filtered genes on autosome *c*. A joint score test yielded concordant gene-pair associations and retained all representative loci (Supplementary Methods; Supplementary Table S2). We used the component-union screen in the primary analysis because it preserved component-specific evidence from the count and detection models.

### Simulation design

#### Hurdle-compatible model-based simulation

We constructed a parametric model-based simulation to evaluate the calibration and power of the hurdle-based co-expression tests. Model parameters were estimated from immature and naïve B cells (B IN), and expression counts were generated using separate distributions for detection (nonzero-expression) probability and positive-count abundance. For cell *c* and gene *g*,

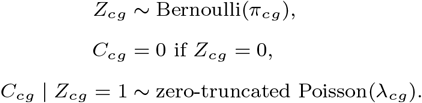

The linear predictors were

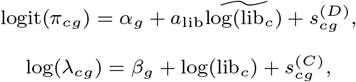

where 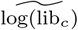 denotes centered log library size. Reference fitting estimated the donor cell-count, library-size, and gene-level detection and positive-count baselines from observed B IN cells. Each replicate comprised 300 donors and 100 genes, and 50 replicates were generated per scenario. In each non-null replicate, 5% of all possible unordered gene pairs were designated as signal pairs, defined as pairs assigned a shared cell-level latent variable to induce co-expression. These pairs were sampled without replacement from the subset in which both genes had reference detection rates and positive-count intensities at or above their respective medians.

For each selected signal pair *p*, we generated an independent cell-level latent variable *h*_*cp*_ *~ N* (0, 1), which was shared by the two genes in that pair. The parameters *γ*_*D*_ and *γ*_*C*_ denote the coefficients of this shared factor in the detection-logit and positive-count log-rate models, respectively; the evaluated scenarios used *γ*_*D*_ ∈ {0, 0.35, 0.60} and *γ*_*C*_ ∈ {0, 0.20, 0.35}. If *P_g_* denotes the selected pairs containing gene *g*, the simulated signal terms were

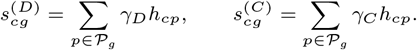

The null setting used (*γ*_*D*_, *γ*_*C*_) = (0, 0). Detection-only settings, labeled zero-only in simulation files and Figure 2, used (0.35, 0) and (0.60, 0) for the lower and higher shared-factor loadings, respectively. Because *h*_*cp*_ has unit variance and enters the detection logits of both genes, these *γ*_*D*_ values correspond to detection odds ratios of exp(0.35) = 1.42 and exp(0.60) = 1.82 per one-standard-deviation increase in the shared factor. Count-only settings used (0, 0.20) and (0, 0.35), corresponding to positive-count Poisson rate ratios of exp(0.20) = 1.22 and exp(0.35) = 1.42, and joint settings combined the respective detection and count loadings as (0.35, 0.20) and (0.60, 0.35). Thus, the lower and higher labels refer to the magnitude of the coefficient linking the shared latent factor to both genes, rather than to a prespecified gene-gene correlation. The data-generating model contained no donor-level covariate effects. In the fitted hurdle models, randomized donor sex was retained as a null adjustment covariate, whereas age, genotype principal components, and PEER factors were constant and therefore omitted automatically. Library size entered the fitted count and detection components as an offset and covariate, respectively.

In each simulation replicate, the two response-predictor directions were analyzed as separate records. Within each direction, *p*_union_ = min(*p*_count_, *p*_detection_), and the test was considered positive when *p*_union_ *< α*, where *α* denotes the nominal per-test significance threshold. We evaluated *α ∈* {0.05, 0.01, 0.005, 0.001}. Results for the count component, detection component, and component-union rule were compared with undirected pseudobulk Pearson and Spearman tests based on donor-level mean expression.

#### Real-data permutation-based simulation

To evaluate calibration and power under realistic single-cell count distributions, we constructed a permutation-based simulation using an observed OneK1K count matrix. Unlike the model-based simulation, this design preserved each gene’s observed marginal count distribution. For the null analysis, we selected 200 genes from B IN cells and independently permuted each gene’s counts across cells. This procedure preserved marginal count distribution, donor labels, and donor-specific cell counts while removing gene-gene dependence and disrupting the original alignment between expression and measured cell- and donor-level covariates.

Counts were permuted across all cells rather than within donors, because within-donor permutation leaves donor-level gene means unchanged and therefore does not eliminate donor-level co-expression (Supplementary Figure S2). Because the permutation disrupted the original alignment with the measured covariates, the hurdle models fitted to the permuted data omitted age, sex, genotype principal components, PEER factors, and library-size terms. Both ordered directions were analyzed separately, as in the model-based simulation. For the power analysis, the permuted matrix served as the background, and shared cell-level effects were introduced into four non-overlapping 20-gene clusters. For cell *c*, gene *g*, and cluster *m* containing gene *g*, the injected effect was

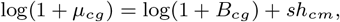

where *B*_*cg*_ denotes the permuted count, *h*_*cm*_ *~ N* (0, 1) is a cell-level latent variable shared by all genes in cluster *m*, and *s* is the log-scale effect coefficient. Counts were then sampled from Poisson(*µ*_*cg*_), with *s* = 0.02, 0.04, or 0.06.

We compared sc-pcQTL with pseudobulk Spearman correlations computed from the same matrices.

### Sliding-window cluster identification

Following previous pcQTL work on bulk data (Lawrence et al., 2026), we identified clusters independently on each chromosome using a greedy sliding-window procedure. Genes were ordered by genomic start position. Starting with windows of 50 adjacent genes, we reduced one gene at a time to a minimum of two. At each size, windows were scanned along the chromosome in one-gene increments. For a window containing *k* genes, we calculated the proportion of its 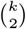 unordered gene pairs that passed the pairwise screen. A window was retained as a cluster if this proportion was at least 70%. Its genes were then marked as assigned, and subsequent windows containing an assigned gene were skipped. This largest-window-first procedure produced non-overlapping clusters, which were then combined across chromosomes.

### Cluster-PC QTL mapping

For each sc-pcQTL-defined cluster, we used principal component analysis (PCA) to summarize the joint expression variation of its constituent genes. PCA was performed separately within each cell type using centered, unscaled SCTransform-corrected counts. Clusters with fewer than two available genes or fewer retained cells than genes were excluded. We retained the minimum number of leading PCs required to explain at least 95% of the cumulative expression variance. Because PC signs are arbitrary, loading-based interpretation focused on the relative directions and absolute magnitudes of gene loadings.

The retained cluster-PC phenotypes were tested for *cis*-pcQTL associations using SAIGE-QTL (Zhou et al., 2024). Models were fitted at the cell level, with donor genotypes and donor-level covariates repeated across cells from the same donor. Age, sex, genotype principal components 1–6, and cell-type-specific PEER factors 1–2 were included as covariates. Variant tests used *±*500-kb *cis* windows around each cluster and were restricted to variants with MAF *≥* 0.05. Cluster-PC phenotypes were analyzed in SAIGE-QTL quantitative-trait mode after inverse-normal-transformation, with variance ratios estimated from pruned markers. Within each cluster-PC phenotype, *cis*-variant *p*-values were adjusted using the Benjamini-Hochberg procedure (Benjamini and Hochberg, 1995); a phenotype was considered significant if at least one variant had *q <* 0.05. In a secondary phenotype-level sensitivity analysis, we combined the *cis*-variant *p*-values for each phenotype using the aggregated Cauchy association test (ACAT) (Liu et al., 2019) and applied Benjamini-Hochberg correction across all successfully tested phenotypes, using *q <* 0.05 as the ACAT-BH threshold.

### OneK1K QTL fine-mapping

We compared cluster-PC pcQTLs with publicly available, cell-type-specific OneK1K single-gene *cis*-eQTL summary statistics generated using SAIGE-QTL (Zhou et al., 2024). Fine-mapping was performed separately for each PCA-selected cluster-PC phenotype with pcQTL summary statistics and for each constituent gene with single-gene eQTL summary statistics from the same cell type. Phenotypes were not required to reach a QTL significance threshold before fine-mapping. Within each cluster, all pcQTL and eQTL phenotypes were fine-mapped across the cluster genomic span extended by 500 kb on each side, matching the pcQTL *cis*-testing region. A common LD matrix was used for all phenotypes in a cluster and was calculated from OneK1K donor genotypes using PLINK (Chang et al., 2015) after restricting variants to MAF *≥* 0.05. All fine-mapping analyses used the hg19 coordinate system.

For each phenotype, we retained SNPs within the fine-mapping interval that were also present in the cluster-specific LD matrix and ordered them to match the matrix. We then aligned the effect alleles in the summary statistics to the LD-matrix allele order. If the two alleles appeared in reverse order, we multiplied 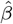 by *™*1 so that the effect estimate referred to the same allele. We excluded SNPs with non-finite effect estimates, standard errors, or *p*-values, as well as SNPs with non-positive standard errors. We applied SuSiE-RSS using `susie_rss` with 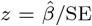, the aligned LD matrix, and the SAIGE-QTL donor sample size (*N* = 982) (Wang et al., 2020). We set the maximum number of effects to *L* = 10, used default prior weights across variants, requested 95% credible sets, and disabled residual-variance estimation. Credible sets were retained only if the minimum absolute pairwise LD correlation among their variants was at least 0.5. Only successful fits containing at least two usable variants and at least one credible set passing this LD criterion were carried forward to colocalization.

### Colocalization with GWAS signals in the FinnGen study

We evaluated colocalization between fine-mapped OneK1K QTL signals and FinnGen R12 SuSiE fine-mapping results (Kurki et al., 2023; FinnGen, 2024). For each cell type and cluster, we compared every successfully fine-mapped single-gene eQTL and cluster-PC pcQTL with traits in the FinnGen study whose fine-mapped GWAS regions overlapped the cluster window. Before colocalization, OneK1K QTL variants were lifted from hg19 to hg38 and matched to variants in the FinnGen study by chromosome, position, and allele compatibility. We ran `coloc.susie` (Giambartolomei et al., 2014; Wallace, 2021) on the shared variants only when at least two variants remained and both SuSiE fits retained a credible set.

To construct signal groups, we linked two fine-mapped signals when a QTL-GWAS or within-cluster eQTL-pcQTL comparison exceeded the selected PPH4 threshold. Signals connected through one or more links were assigned to the same group, and only groups containing at least one GWAS signal were retained. eQTL-only groups contained eQTL but no pcQTL signals, shared groups contained both QTL types, and pcQTL-specific groups contained at least one pcQTL signal but no colocalized eQTL from a constituent gene. The primary threshold was PPH4 > 0.75; thresholds of 0.70 and 0.80 were evaluated in sensitivity analyses. When counting unique GWAS hits, repeated colocalizations involving the same GWAS phenotype and GWAS lead variant were counted only once, regardless of cell type, cluster, or QTL phenotype.

## Discussion

sc-pcQTL combines hurdle-based co-expression screening with cluster-PC phenotypes to summarize local multi-gene expression variation. By modeling detection and positive-count abundance separately, the framework accommodates expression sparsity and library-size variation. In simulations, the individual hurdle components were well calibrated and substantially more powerful than pseudobulk correlation tests, particularly for detection-linked and joint effects.

Using the same fine-mapping, colocalization framework for pcQTLs and eQTLs, we identified pcQTL signals that colocalized with GWAS loci but had no colocalized single-gene eQTL among their constituent genes. These findings indicate that aggregating coordinated expression across neighboring genes can complement conventional single-gene analysis and improve detection of genetically regulated, trait-relevant expression patterns.

By analyzing defined immune cell types, sc-pcQTL can resolve multi-gene regulatory signals that may be diluted by bulk or pseudobulk averaging. In the *GIMAP* example and pseudobulk mixture simulation, the cluster-PC colocalization signal was strongest when the relevant cell type was analyzed separately. Although the same seven-gene module was identified in several immune cell types, colocalization with GWAS of lymphocyte count exceeded the primary threshold only for the cluster-PC phenotype in CD8 NC cells. This example illustrates the potential of sc-pcQTL to identify cell-type-specific multi-gene regulatory signals.

This study has several limitations. First, it focused on European-ancestry OneK1K immune-cell data, the portability of the findings across cohorts, ancestries, tissues, and disease-relevant cell states remains to be established. Second, comparisons across cell types were also influenced by differences in donor and cell numbers, expression sparsity, and the genes eligible for testing. Finally, the pairwise hurdle model provided a computationally feasible approach to genome-wide feature construction across more than one million cells. Although within-donor dependence was not explicitly modeled during screening, a sensitivity analysis found no consistent increase in rejection rates at the evaluated significance levels (Supplementary Figure S3). This conclusion is limited to the within-donor dependence patterns represented in this OneK1K-based simulation.

Future applications to independent datasets spanning diverse ancestries, tissues, and cell types will help establish the generalizability of sc-pcQTL across study settings. Integrating chromatin data, perturbation experiments, and functional assays could further strengthen the biological interpretation of specific pcQTL modules. Extending the framework beyond local *cis* regulation to trans effects and broader gene-cluster discovery may reveal additional coordinated regulatory programs.

## Ethics approval and consent to participate

This secondary analysis used previously generated, de-identified OneK1K resources; recruitment, consent, ethics approval, and primary data generation were described in the original study (Yazar et al., 2022).

## Data availability

OneK1K expression data are available through GEO under accession GSE196830 (Gene Expression Omnibus, 2022); access to the associated genotype and covariate data is described in the original study (Yazar et al., 2022). Public OneK1K eQTL summary statistics using SAIGE-QTL are available from Zenodo (https://doi.org/10.5281/zenodo.10884040) (Zhou et al., 2024). FinnGen R12 GWAS summary statistics and SuSiE fine-mapping results were obtained from the FinnGen public data release (FinnGen, 2024), and GWAS summary statistics for study GCST90002096 were obtained from the NHGRI-EBI GWAS Catalog (NHGRI-EBI GWAS Catalog, 2020). The summary result tables generated in this study are publicly available on Zenodo (DOI: https://doi.org/10.5281/zenodo.21222687)

## Code availability

The sc-pcQTL software is openly available at https://github.com/ZhouLabGenetics/sc-pcQTL. The analysis workflow and figure-generation code used in this study are openly available at https://github.com/ZhouLabGenetics/sc-pcQTL_code.

## Supplementary information

Supplementary material includes figures, diagnostics, and representative-locus plots.

## Funding

W.Z. was supported by the National Human Genome Research Institute of the National Institutes of Health under award numbers R00HG012222 and R01HG014518. M.C. was supported by the Weissman Family MGH Research Scholar Award. M.C. and Y.H. were further supported by the Novo Nordisk Foundation (NNF21SA0072102), UM1DK126185, RC2 DK144819-01, P30 DK040561.

## Acknowledgements

We thank Dr. Joseph Powell for facilitating access to the OneK1K dataset. We also thank the participants of the OneK1K and FinnGen studies.

## Use of AI tools

Anthropic Claude Code and OpenAI Codex assisted with code editing, language editing, formatting checks, and submission-readiness notes. The authors reviewed all outputs and are responsible for the final content.

## Author contributions

J.Z.: Conceptualization, Methodology, Software, Formal analysis, Data curation, Visualization, Writing-original draft. Y.H.: Visualization, Downstream analysis, Writing-review and editing. M.C.: Resources, Writing-review and editing. M.K.: Conceptualization, Methodology, Supervision, Resources, Writing-review and editing. W.Z.: Conceptualization, Methodology, Supervision, Project administration, Resources, Writing-review and editing.

## Conflict of interest

None declared.

1 Note: in this study, pseudobulk expression was calculated as the mean count of each gene across all cells from the same donor within a cell type; gene-pair correlations were then evaluated across donors.

## Notes

https://github.com/ZhouLabGenetics/sc-pcQTL

https://github.com/ZhouLabGenetics/sc-pcQTL_code

https://doi.org/10.5281/zenodo.21222687

### Competing Interest Statement

The authors have declared no competing interest.

## References

Y. Benjamini and Y. Hochberg. Controlling the false discovery rate: a practical and powerful approach to multiple testing. Journal of the Royal Statistical Society: Series B, 57(1):289–300, 1995. doi: 10.1111/j.2517-6161.1995.tb02031.x.

B. K. Bulik-Sullivan et al. LD Score regression distinguishes confounding from polygenicity in genome-wide association studies. Nature Genetics, 47(3):291–295, 2015. doi: 10.1038/ng.3211.

C. C. Chang, C. C. Chow, L. C. Tellier, S. Vattikuti, S. M. Purcell, and J. J. Lee. Second-generation PLINK: rising to the challenge of larger and richer datasets. GigaScience, 4:7, 2015. doi: 10.1186/s13742-015-0047-8.

A. S. E. Cuomo et al. Single-cell RNA-sequencing of differentiating iPS cells reveals dynamic genetic effects on gene expression. Nature Communications, 11:810, 2020. doi: 10.1038/s41467-020-14457-z.

FinnGen. FinnGen Documentation of R12 release. FinnGen Public Documentation, 2024. URL https://finngen.gitbook.io/documentation/r12. [dataset].

H. K. Finucane et al. Partitioning heritability by functional annotation using genome-wide association summary statistics. Nature Genetics, 47(11):1228–1235, 2015. doi: 10.1038/ng.3404.

S. Gazal et al. Linkage disequilibrium-dependent architecture of human complex traits shows action of negative selection. Nature Genetics, 49(10):1421–1427, 2017. doi: 10.1038/ng.3954.

Gene Expression Omnibus. Single-cell eQTL mapping identifies cell type specific genetic control of autoimmune disease. GEO Series GSE196830, 2022. URL https://www.ncbi.nlm.nih.gov/geo/query/acc.cgi?acc=GSE196830. [dataset].

A. T. Ghanbarian and L. D. Hurst. Neighboring genes show correlated evolution in gene expression. Molecular Biology and Evolution, 32(7):1748–1766, 2015. doi: 10.1093/molbev/msv053.

C. Giambartolomei et al. Bayesian test for colocalisation between pairs of genetic association studies using summary statistics. PLOS Genetics, 10(5):e1004383, 2014. doi: 10.1371/journal.pgen.1004383.

C. Hafemeister and R. Satija. Normalization and variance stabilization of single-cell RNA-seq data using regularized negative binomial regression. Genome Biology, 20(1):296, 2019. doi: 10.1186/s13059-019-1874-1.

J. Hu, J. N. Weber, L. E. Fuess, N. C. Steinel, D. I. Bolnick, and M. Wang. A spectral framework to map QTLs affecting joint differential networks of gene co-expression. PLOS Computational Biology, 21(4):e1012953, 2025. doi: 10.1371/journal.pcbi.1012953.

Y. Huang, M. J. A. M. Verstegen, S. J. D. Tjalsma, et al. Two unrelated distal genes activated by a shared enhancer benefit from localizing inside the same small topological domain. Genes & Development, 39(5–6):348–363, 2025. doi: 10.1101/gad.352235.124.

M. Kanai, T. M. Delorey, J. Honkanen, R. S. Rodosthenous, J. Juvila, S. Murphy, et al. Population-scale multiome immune cell atlas reveals complex disease drivers. medRxiv, 2025. doi: 10.1101/2025.11.25.25340489. Preprint.

D. Kaptijn, C. Losert, M. Korshevniuk, et al. Disease-associated variants are enriched for altering cell-type-specific gene co-expression relationships. bioRxiv, 2025. doi: 10.1101/2025.09.06.674678. Preprint.

S. Khetan, B. S. Carroll, and M. L. Bulyk. Multiple overlapping binding sites determine transcription factor occupancy. Nature, 646:1001–1011, 2025. doi: 10.1038/s41586-025-09472-3.

M. I. Kurki et al. FinnGen provides genetic insights from a well-phenotyped isolated population. Nature, 613:508–518, 2023. doi: 10.1038/s41586-022-05473-8.

K. A. Lawrence, T. Gjorgjieva, D. Nachun, and S. B. Montgomery. Focus on single-gene effects limits discovery and interpretation of complex-trait-associated variants. The American Journal of Human Genetics, 113(4):842–851, 2026. doi: 10.1016/j.ajhg.2026.02.022.

S. Li, K. T. Schmid, D. H. de Vries, et al. Identification of genetic variants that impact gene co-expression relationships using large-scale single-cell data. Genome Biology, 24:80, 2023. doi: 10.1186/s13059-023-02897-x.

M.-A. Limoges, M. Cloutier, M. Nandi, S. Ilangumaran, and S. Ramanathan. The GIMAP family proteins: an incomplete puzzle. Frontiers in Immunology, 12:679739, 2021. doi: 10.3389/fimmu.2021.679739.

H. Liu, A. Abedini, E. Ha, et al. Kidney multiome-based genetic scorecard reveals convergent coding and regulatory variants. Science, 387(6734):eadp4753, 2025. doi: 10.1126/science.adp4753.

Y. Liu, S. Chen, Z. Li, A. C. Morrison, E. Boerwinkle, and X. Lin. ACAT: A fast and powerful p value combination method for rare-variant analysis in sequencing studies. The American Journal of Human Genetics, 104(3):410–421, 2019. doi: 10.1016/j.ajhg.2019.01.002.

J. Mullahy. Specification and testing of some modified count data models. Journal of Econometrics, 33(3):341–365, 1986. doi: 10.1016/0304-4076(86)90002-3.

NHGRI-EBI GWAS Catalog. GWAS Catalog study GCST90002096: CD11b on basophil. NHGRI-EBI GWAS Catalog, 2020. URL https://www.ebi.ac.uk/gwas/studies/GCST90002096. [dataset].

J. C. Pascall, L. M. C. Webb, E.-L. Eskelinen, S. Innocentin, N. Attaf-Bouabdallah, and G. W. Butcher. GIMAP6 is required for T cell maintenance and efficient autophagy in mice. PLOS ONE, 13(5):e0196504, 2018. doi: 10.1371/journal.pone.0196504.

K. Pearson. On lines and planes of closest fit to systems of points in space. The London, Edinburgh, and Dublin Philosophical Magazine and Journal of Science, 2(11):559–572, 1901. doi: 10.1080/14786440109462720.

A. L. Price, N. J. Patterson, R. M. Plenge, M. E. Weinblatt, N. A. Shadick, and D. Reich. Principal components analysis corrects for stratification in genome-wide association studies. Nature Genetics, 38:904–909, 2006. doi: 10.1038/ng1847.

D. M. Ribeiro et al. The molecular basis, genetic control and pleiotropic effects of local gene co-expression. Nature Communications, 12:4842, 2021. doi: 10.1038/s41467-021-25129-x.

A. Saha and A. Battle. False positives in trans-eQTL and co-expression analyses arising from RNA-sequencing alignment errors. F1000Research, 7:1860, 2018. doi: 10.12688/f1000research.17145.2.

A. Saunders, L. M. C. Webb, M. L. Janas, A. Hutchings, J. Pascall, C. Carter, et al. Putative GTPase GIMAP1 is critical for the development of mature B and T lymphocytes. Blood, 115(16): 3249–3257, 2010. doi: 10.1182/blood-2009-08-237586.

S. Schoenfelder, T. Sexton, L. Chakalova, et al. Preferential associations between co-regulated genes reveal a transcriptional interactome in erythroid cells. Nature Genetics, 42(1):53–61, 2010. doi: 10.1038/ng.496.

R. D. Schulteis, H. Chu, X. Dai, Y. Chen, B. Edwards, D. Haribhai, et al. Impaired survival of peripheral T cells, disrupted NK/NKT cell development, and liver failure in mice lacking Gimap5. Blood, 112(13):4905–4914, 2008. doi: 10.1182/blood-2008-03-146555.

O. Stegle, L. Parts, M. Piipari, J. Winn, and R. Durbin. Using probabilistic estimation of expression residuals (PEER) to obtain increased power and interpretability of gene expression analyses. Nature Protocols, 7:500–507, 2012. doi: 10.1038/nprot.2011.457.

C. Su et al. Cell-type-specific co-expression inference from single cell RNA-sequencing data. Nature Communications, 14:4846, 2023. doi: 10.1038/s41467-023-40503-7.

U. Vósa et al. Large-scale cis-and trans-eQTL analyses identify thousands of genetic loci and polygenic scores that regulate blood gene expression. Nature Genetics, 53:1300–1310, 2021. doi: 10.1038/s41588-021-00913-z.

M. G. P. van der Wijst et al. Single-cell RNA sequencing identifies cell-type-specific cis-eQTLs and co-expression QTLs. Nature Genetics, 50(4):493–497, 2018. doi: 10.1038/s41588-018-0089-9.

C. Wallace. A more accurate method for colocalisation analysis allowing for multiple causal variants. PLoS Genetics, 17(9): e1009440, 2021. doi: 10.1371/journal.pgen.1009440.

G. Wang, A. Sarkar, P. Carbonetto, and M. Stephens. A simple new approach to variable selection in regression, with application to genetic fine mapping. Journal of the Royal Statistical Society: Series B, 82(5):1273–1300, 2020. doi: 10.1111/rssb.12388.

C. A. R. Warmerdam et al. Trans-eQTLs reveal the architecture of human gene regulatory networks. medRxiv, 2026. doi: 10.64898/2026.02.04.26343575. Preprint.

S. Yazar et al. Single-cell eQTL mapping identifies cell type-specific genetic control of autoimmune disease. Science, 376 (6589):eabf3041, 2022. doi: 10.1126/science.abf3041.

W. Zhou et al. Efficient and accurate mixed model association tool for single-cell eQTL analysis. medRxiv, 2024. doi: 10.1101/2024.05.15.24307317. URL https://www.medrxiv.org/content/10.1101/2024.05.15.24307317v1. Preprint.

Z. Zhu et al. Integration of summary data from GWAS and eQTL studies predicts complex trait gene targets. Nature Genetics, 48(5):481–487, 2016. doi: 10.1038/ng.3538.

K. D. Zimmerman, M. A. Espeland, and C. D. Langefeld. A practical solution to pseudoreplication bias in single-cell studies. Nature Communications, 12:738, 2021. doi: 10.1038/s41467-021-21038-1.

