## Supplementary Material for "sc-pcQTL: hurdle-based co-expression modeling for multi-gene QTL mapping in single-cell RNA-seq data"

### Supplementary analyses

#### Cell-type eligibility

Four cell types had fewer than 10,000 annotated cells and were excluded before pairwise screening: CD4 SOX4 T cells (CD4_SOX4; 4,065 cells), dendritic cells (DC; 8,690 cells), NK recruiting cells (NK_R; 9,677 cells), and plasma cells (Plasma; 3,625 cells). Cell counts and the corresponding exclusion reasons are summarized in Supplementary Table S1.

**Supplementary Table S1:** Cell-type eligibility for primary analyses. Cell types with at least 10,000 annotated cells were included before pairwise screening, and the resulting 10-cell-type set was used throughout the primary analyses.

| OneK1K label | Annotation | Cells | Status | Exclusion reason |
|---|---|---|---|---|
| B_IN | Immature/naïve B | 82,068 | Included | None |
| B_Mem | Memory B | 48,023 | Included | None |
| CD4_ET | CD4 effector T | 61,786 | Included | None |
| CD4_NC | CD4 naïve T | 463,528 | Included | None |
| CD4_SOX4 | CD4 SOX4 T | 4,065 | Excluded | Fewer than 10,000 cells |
| CD8_ET | CD8 effector T | 205,077 | Included | None |
| CD8_NC | CD8 naïve T | 133,482 | Included | None |
| CD8_S100B | CD8 S100B T | 34,528 | Included | None |
| DC | Dendritic | 8,690 | Excluded | Fewer than 10,000 cells |
| Mono_C | Classical monocyte | 38,233 | Included | None |
| Mono_NC | Nonclassical monocyte | 15,166 | Included | None |
| NK | Natural killer | 159,820 | Included | None |
| NK_R | NK recruiting | 9,677 | Excluded | Fewer than 10,000 cells |
| Plasma | Plasma | 3,625 | Excluded | Fewer than 10,000 cells |

#### Exploratory observed-data diagnostic of sc-pcQTL hurdle and pseudobulk co-expression

We performed a diagnostic analysis in immature and naïve B cells (B_IN) from OneK1K to illustrate pairwise patterns captured differently by the sc-pcQTL hurdle model and donor-level pseudobulk Spearman correlation.

Three representative gene pairs illustrate distinct sources of disagreement (Supplementary Figure S1). For *ATP13A3-RPL35A*, the exploratory single-cell analysis revealed differences in both detection probability and positive counts among cells with nonzero expression. Averaging all cells within each donor eliminated both sources of variation, yielding no detectable pseudobulk correlation.

For *MIF-MTFP1*, the association was driven primarily by cell-level detection and was therefore detected mainly by the detection component. Pseudobulk averaging combined detection and positive-count information into a single mean, obscuring this pattern.

*ATG7-HACL1* illustrated the converse pattern. Both genes were expressed in fewer than 3% of cells, and neither hurdle component detected a cell-level association. Their pseudobulk values nevertheless formed near-linear bands and produced a significant Spearman correlation. This arose because both gene means shared the same donor-specific cell-count denominator, creating a common source of variation.

Together, these examples illustrate two complementary limitations of pseudobulk correlation for sparse single-cell data. Pseudobulk averaging can obscure cell-level associations carried by detection and positive counts, whereas a shared donor-specific cell-count denominator can induce apparent correlations between ultra-sparse genes. By modeling detection and positive counts separately at the cell level, the sc-pcQTL hurdle approach retained both forms of pairwise information in this diagnostic analysis.

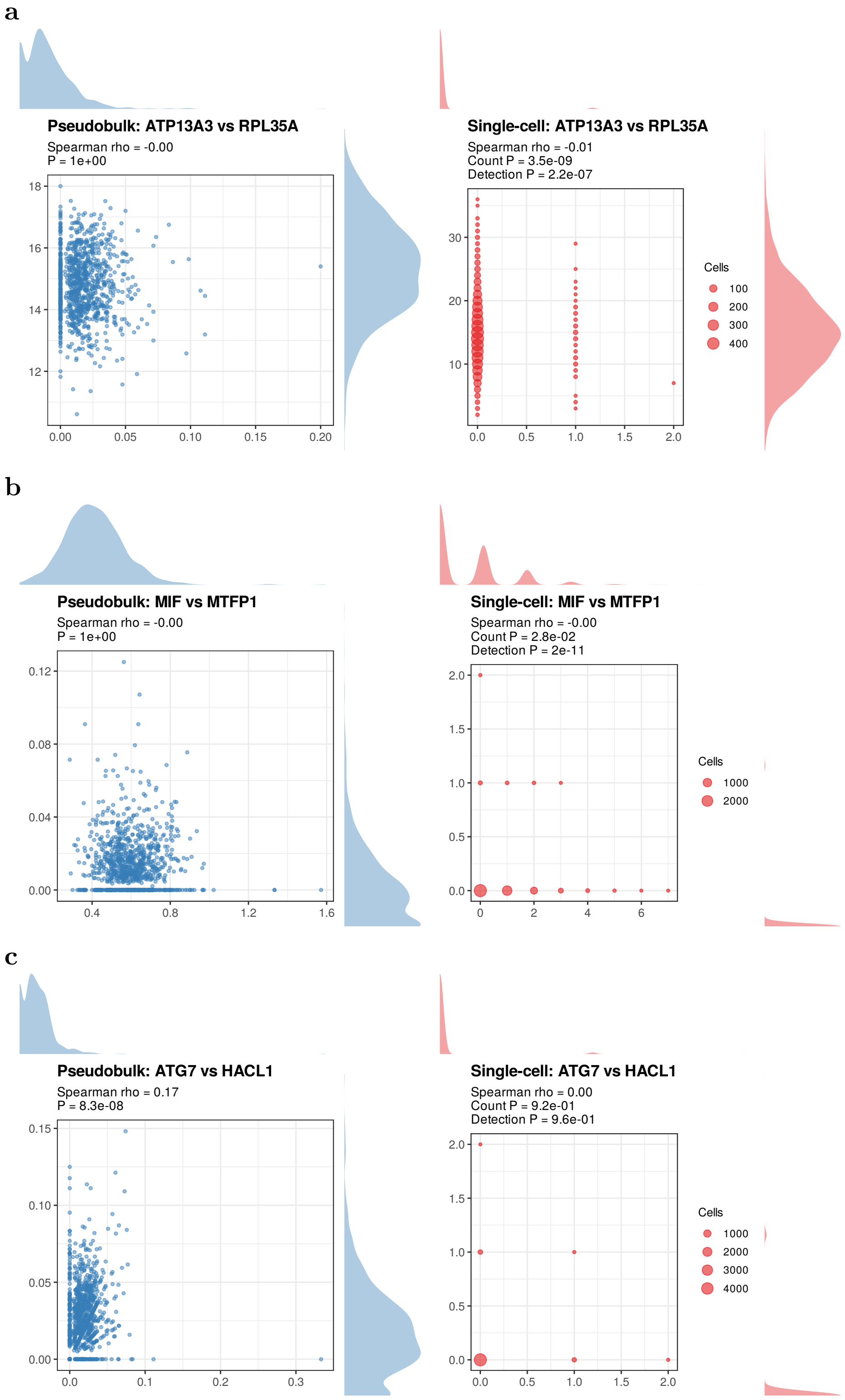


**Supplementary Figure S1:** Exploratory observed-data diagnostic of pairwise disagreement between pseudobulk Spearman and the sc-pcQTL hurdle analysis in B_IN cells. (a) *ATP13A3-RPL35A*, for which cell-level detection and positive-count differences were both removed by pseudobulk averaging. (b) *MIF-MTFP1*, for which the association was concentrated in the detection pattern and was detected primarily by the detection component. (c) *ATG7-HACL1*, a pseudobulk-only association induced by the shared donor-specific cell-count denominator: the two ultra-sparse gene means formed near-linear bands despite no evidence from either hurdle component. Point size in the single-cell panels represents the number of sampled cells at each count combination. These panels were generated independently of the primary pairwise screen and were not used in cluster construction or downstream QTL analyses.

### Diagnostic comparison of global and within-donor permutation

We repeated the real-data null simulation using the same analysis settings as the primary simulation, except that counts were permuted within each donor rather than across all cells. Compared with global gene-wise permutation, within-donor permutation yielded substantially higher empirical rejection rates for both hurdle components, the component-union rule, and pseudobulk Spearman correlation. These findings show that within-donor permutation retains substantial donor-level co-expression, supporting the use of global gene-wise permutation for the primary null analysis.

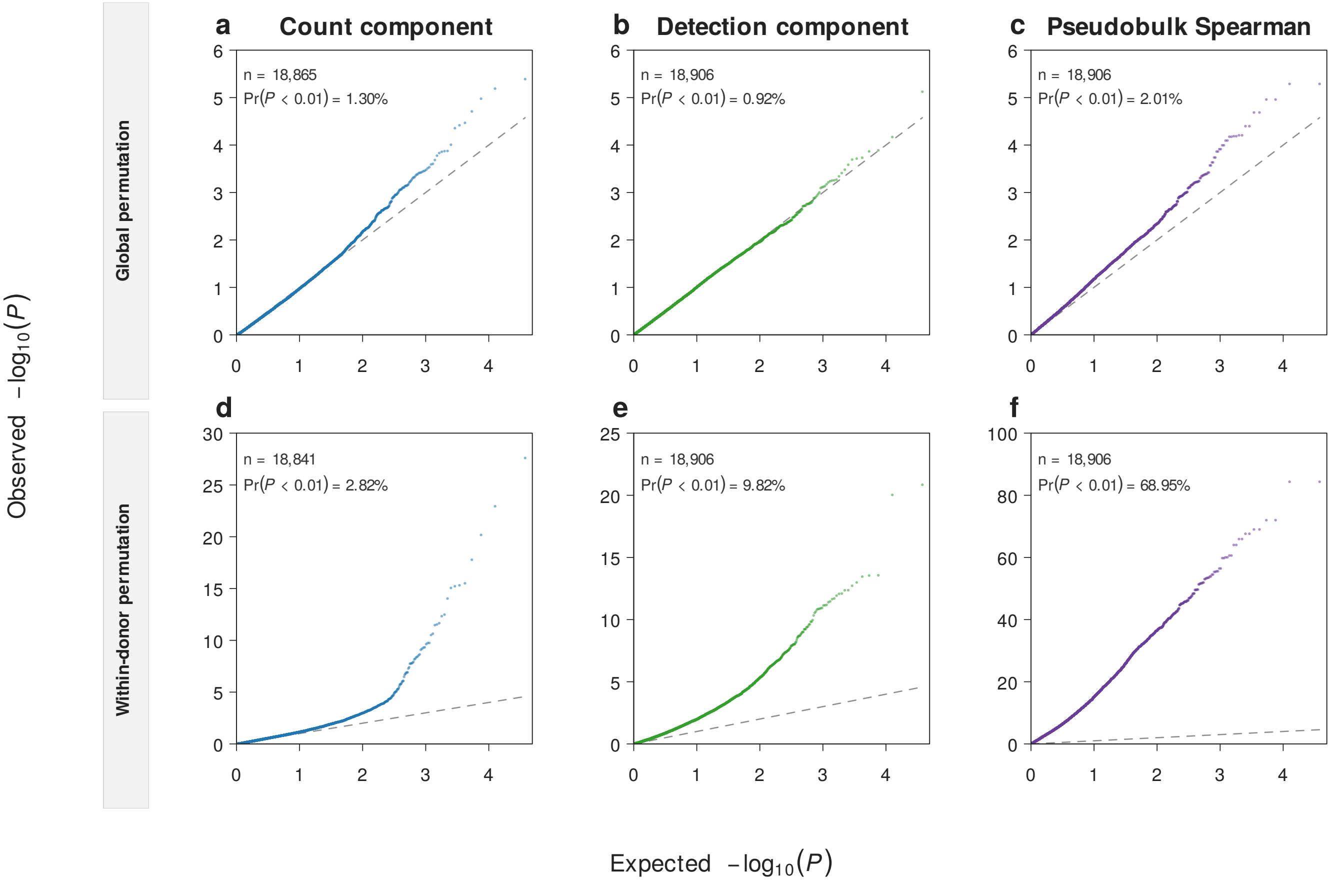


**Supplementary Figure S2:** Within-donor permutation retains donor-level co-expression structure. Quantile-quantile plots compare global gene-wise permutation (a–c) with within-donor gene-wise permutation (d–f) for the Poisson count component, binomial detection component, and donor-level pseudobulk Spearman test. Each point represents one ordered gene-pair test among pairs passing the 1% nonzero-cell cutoff; $n$ denotes the number of finite $p$-values. Dashed lines show the uniform-null expectation, and panel annotations report the fraction of tests with $P < 0.01$. Global permutation disrupted cell- and donor-level gene-gene association, whereas within-donor permutation preserved every donor-gene mean. The within-donor fractions are therefore apparent positive fractions and not valid empirical false-positive rates.

### OneK1K-calibrated within-donor dependence sensitivity analysis

We assessed whether within-donor cell dependence affected null calibration. For each of the 10 primary cell types, one 32-gene panel was sampled and then fixed across simulations. Four genes represented each sparsity level, defined by nonzero-expression frequencies of 1–2%, 2–5%, 5–10%, 10–20%, 20–40%, 40–70%, 70–90%, or 90–99%. Let $N_s$ be the number of eligible genes in level $s$ and $m_s = 4$ the number in the panel. A panel gene in level $s$ received weight $w_g = N_s/m_s$. In each replicate, the first gene was sampled with probability proportional to $w_g$. The second was sampled from the remaining 31 genes after renormalizing the weights. Consequently, the first gene came from level $s$ with probability $N_s/\sum_t N_t$. Sequential sampling approximately retained this single-gene level distribution across both pair members, but did not reproduce the complete eligible-gene pair distribution.

Detection dependence was estimated from a binomial-logit random-intercept model fitted to a balanced sample of at most 50 cells per donor:

$$\operatorname{logit} \Pr(D_{cdg} = 1) = \boldsymbol{x}_{cd}^{(D)\top} \boldsymbol{\gamma}_g + b_{dg}, \qquad b_{dg} \sim N(0, \sigma^2_{D,g}),$$

where $D_{cdg}$ indicates nonzero expression and $\boldsymbol{x}_{cd}^{(D)}$ contains age, sex, genotype principal components 1–6, PEER factors 1–2, and standardized log library size. We used the estimated random-intercept standard deviation $\hat{\sigma}_{D,g}$. The fixed-effect coefficients $\hat{\boldsymbol{\gamma}}_g$ used for simulation were estimated from the same covariates across all cells. Estimates were replaced when the required model fits failed, lacked convergence or a positive-definite Hessian, or had a nonfinite or greater-than-2 standard error for the log standard deviation. Replacement values were the median reliable $\hat{\sigma}_{D,g}$ in the same sparsity level, with the overall median used when no level-specific estimate was available.

Positive-count dependence was estimated among cells with $Y_{cdg} > 0$. The fitted underlying Poisson rate was

$$\lambda_{cdg} = \exp\left\{\log L_{cd} + \boldsymbol{x}_{cd}^{(C)\top} \hat{\boldsymbol{\beta}}_g\right\},$$

where $L_{cd}$ is library size and $\boldsymbol{x}_{cd}^{(C)}$ contains age, sex, genotype principal components 1–6, and PEER factors 1–2. The free intercept in $\hat{\boldsymbol{\beta}}_g$ ensured that the average fitted zero-truncated Poisson conditional mean matched the observed mean among positive cells. We then calculated Pearson residuals

$$r_{cdg} = \frac{Y_{cdg} - \mu(\lambda_{cdg})}{\sqrt{V(\lambda_{cdg})}}, \qquad \mu(\lambda) = \frac{\lambda}{1 - \exp(-\lambda)}, \qquad V(\lambda) = \mu(\lambda)\{1 + \lambda - \mu(\lambda)\}.$$

For variance-component estimation, each donor contributed at most 50 positive cells. Let $K$ be the number of represented donors, $n_d$ the number of retained positive cells from donor $d$, $n = \sum_d n_d$, $\bar{r}_d$ the donor-specific residual mean, and $\bar{r}$ the overall residual mean. We calculated

$$SS_B = \sum_{d=1}^{K} n_d (\bar{r}_d - \bar{r})^2, \qquad SS_W = \sum_{d=1}^{K} \sum_{c=1}^{n_d} (r_{cdg} - \bar{r}_d)^2,$$

$$MS_B = \frac{SS_B}{K-1}, \qquad MS_W = \frac{SS_W}{n-K}, \qquad n_0 = \frac{n - \sum_{d=1}^{K} n_d^2/n}{K-1}.$$

The method-of-moments variance estimates were

$$\hat{\sigma}^2_{b,g} = \max\left\{\frac{MS_B - MS_W}{n_0}, 0\right\}, \qquad \hat{\sigma}^2_{e,g} = \max\{MS_W, 0\},$$

and the observed count-scale intraclass correlation was $\hat{\rho}_g^{\text{obs}} = \hat{\sigma}_{b,g}^2/(\hat{\sigma}_{b,g}^2 + \hat{\sigma}_{e,g}^2)$.

Because discretization attenuates count-scale correlation, $\hat{\rho}_g^{\text{obs}}$ was not used directly as the Gaussian-copula parameter. For each gene, 20 recovery simulations were run at a known latent correlation $\rho_0 = 0.05$ using the same fitted means and donor layout. Let $\hat{\rho}_{gr}^{\text{rec}}$ denote the count-scale ICC recovered in replicate $r$. With $a_g = \text{median}_r(\hat{\rho}_{gr}^{\text{rec}})/\rho_0$, the calibrated latent correlation was

$$\hat{\rho}_g^{\text{lat}} = \min\left\{0.95, \max\left(0, \frac{\hat{\rho}_g^{\text{obs}}}{a_g}\right)\right\}.$$

A direct estimate was considered unstable if the observed correlation or recovery summaries were nonfinite, fewer than 10 recovery estimates were finite, $a_g < 0.5$, the recovery standard deviation exceeded 0.025, fewer than 30 donors had repeated positive cells, or fewer than 200 within-donor positive-cell pairs were available. Unstable estimates were replaced by the 75th percentile of reliable $\hat{\rho}_g^{\text{lat}}$ values in the same positive-count-mean bin $[1, 1.10)$, $[1.10, 1.25)$, $[1.25, 1.50)$, $[1.50, 2)$, or $[2, \infty)$; the overall 75th percentile was the fallback.

For scenario $s$, detection was generated from

$$\operatorname{logit} \Pr(D_{cdg}^{(s)} = 1) = \boldsymbol{x}_{cd}^{(D)\top}\hat{\boldsymbol{\gamma}}_g + \delta_{D,g}^{(s)} + \sigma_{D,g}^{(s)} u_{dg}.$$

Positive counts were generated using

$$z_{cdg}^{(s)} = \sqrt{\rho_{C,g}^{(s)}}\, v_{dg} + \sqrt{1 - \rho_{C,g}^{(s)}}\, \epsilon_{cdg}, \qquad Y_{cdg}^{(s)} = F^{-1}_{\text{ZTP}(\lambda_{cdg}^{(s)})}\left\{\Phi(z_{cdg}^{(s)})\right\} \quad \text{when } D_{cdg}^{(s)} = 1,$$

where $\lambda_{cdg}^{(s)} = \exp\{\log L_{cd} + \boldsymbol{x}_{cd}^{(C)\top}\hat{\boldsymbol{\beta}}_g + \delta_{C,g}^{(s)}\}$. The inverse-CDF step retained the cell-specific zero-truncated Poisson margin defined by $\lambda_{cdg}^{(s)}$. The donor effects $u_{dg}$ and $v_{dg}$, and the cell-level residual $\epsilon_{cdg}$, were independent standard-normal variables in the primary scenario. We numerically solved $\delta_{D,g}^{(s)}$ to match the observed nonzero-expression frequency. We similarly solved $\delta_{C,g}^{(s)}$ so that the average conditional mean $E(Y_{cdg}^{(s)} \mid Y_{cdg}^{(s)} > 0)$ matched the observed positive-count mean.

For each cell type, we generated 10,000 null gene-pair replicates. Each replicate independently sampled one pair from the fixed panel and generated matched dependence and no-donor datasets. The dependence scenario used $(\sigma_{D,g}^{(s)}, \rho_{C,g}^{(s)}) = (\hat{\sigma}_{D,g}, \hat{\rho}_g^{\text{lat}})$, whereas the no-donor scenario set both parameters to zero. For each gene, the matched datasets used the same donor-cell structure, covariates, Bernoulli uniforms, and normal draws. Across the two genes, the Bernoulli uniforms, cell-level Gaussian residuals, and donor-level Gaussian effects were generated independently. The genes shared the fixed donor-cell layout, covariates, and library sizes. The same covariates and library-size adjustments were included in the fitted hurdle models. Both response-predictor directions were analyzed with the primary two-component screen.

Across the 10 cell types, introducing donor dependence produced small changes in null rejection rates with no consistent direction (Supplementary Figure S3). Pooled differences were close to zero, and no cell-type comparison remained significant after Benjamini-Hochberg correction. Monte Carlo intervals reflected simulation uncertainty conditional on the estimated parameters and did not include parameter-estimation uncertainty. Within the OneK1K-based dependence patterns examined, within-donor dependence did not systematically increase null rejection rates.

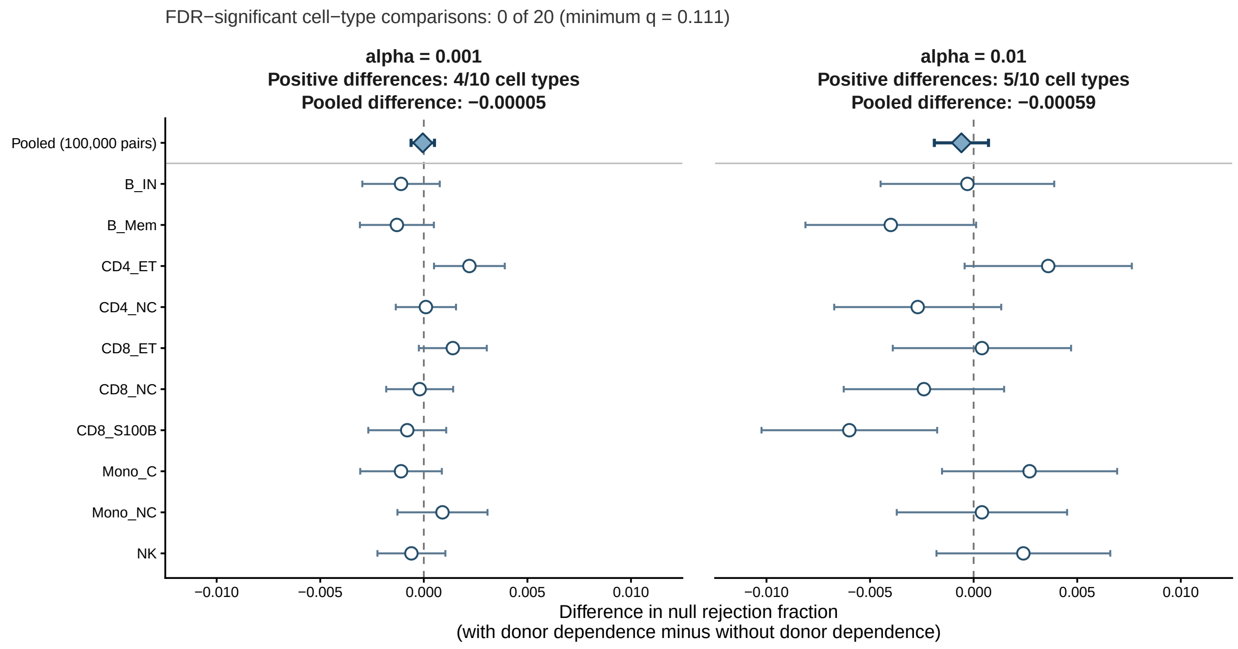


**Supplementary Figure S3:** Within-donor dependence sensitivity analysis. Paired change in the complete-screen null rejection fraction after introducing within-donor dependence estimated from OneK1K. The complete screen uses the smallest of the four $p$-values from the two hurdle components fitted in both response-predictor directions. Each cell type contributed 10,000 matched unordered null pairs per scenario. Open circles show cell-type estimates; diamonds pool all 100,000 matched pairs at each threshold. Facet headers report the number of cell types with a positive difference and the pooled difference. Error bars are unadjusted 95% Monte Carlo intervals conditional on the fitted dependence parameters. Two cell-type intervals excluded zero, but the differences had opposite signs and none of the 20 cell-type comparisons passed Benjamini-Hochberg correction. The pooled differences were $-0.00005$ at $\alpha = 0.001$ and $-0.00059$ at $\alpha = 0.01$, with both intervals crossing zero.

### Count-model calibration under permutation and controlled null simulations

Using the globally permuted OneK1K matrix from the primary null simulation, we refitted each eligible ordered gene-pair test with either a Poisson or Negative-Binomial count component while retaining the same binomial detection component. At $\alpha = 0.01$, the Poisson and Negative-Binomial Wald tests had empirical rejection rates of 0.0130 and 0.0381, respectively, showing that Poisson was better calibrated in this empirical null (Supplementary Figure S4).

To evaluate distribution-dependent calibration, we simulated 20,000 null gene pairs by generating the two genes independently in 20,000 cells, with a 12% detection probability for each gene. Conditional on detection, counts followed one of three regimes: zero-truncated Poisson with mean 2; shifted Poisson, using $1 + \text{Poisson}(1.0)$ for the predictor and $1 + \text{Poisson}(0.6)$ for the response; or zero-truncated Negative-Binomial with mean 2 and dispersion $\theta = 2$. Each simulated pair was fitted with both count models.

Under zero-truncated Poisson counts, both models were calibrated at $\alpha = 0.01$, with rejection rates of 0.00975. Under shifted-Poisson counts, the Negative-Binomial test became anti-conservative in the tail: at $\alpha = 0.001$, its rejection rate was 0.00320, compared with 0.00025 for Poisson, and its median dispersion estimate was $9.25 \times 10^4$. Under overdispersed counts, the pattern reversed: at $\alpha = 0.01$, rejection rates were 0.0311 for Poisson and 0.00935 for Negative-Binomial, and the median estimated dispersion was 2.01 (Supplementary Figure S5a,b). Thus, Poisson was better suited to the SCTransform-corrected OneK1K counts, whereas Negative-Binomial regression remained appropriate when positive counts were overdispersed.

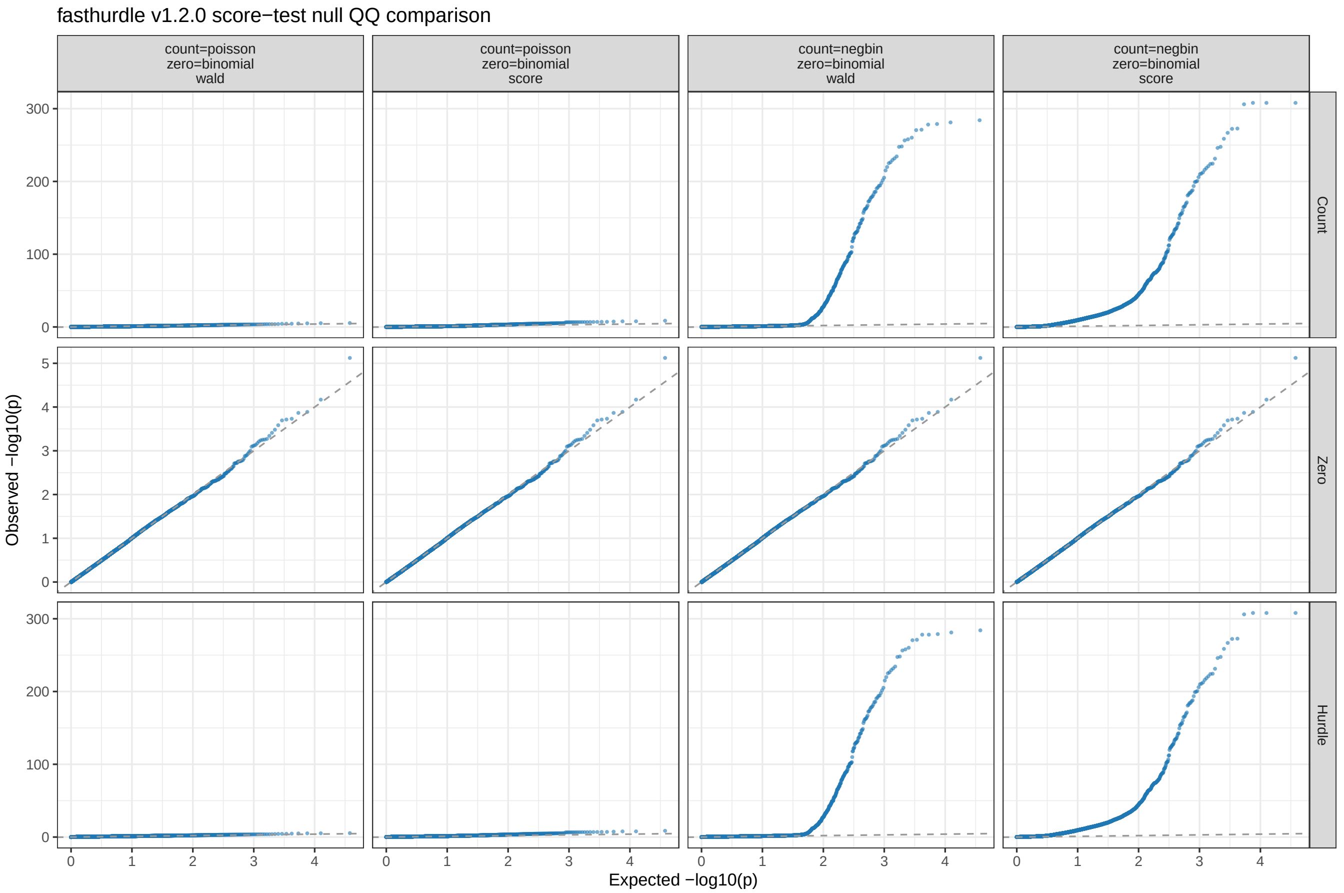


**Supplementary Figure S4:** Count-model calibration in the globally permuted OneK1K matrix. Quantile-quantile plots show the count component, detection component (labeled “Zero” in the original analysis output), and component-union minimum under Poisson Wald, Poisson score, Negative-Binomial Wald, and Negative-Binomial score settings in *fasthurdle* v1.2.0. All settings used the same 9,453 eligible unordered gene pairs evaluated in both ordered directions, a binomial detection model, and no covariates or library-size term. In the score-test columns, the score test was applied only to the count component. Dashed lines show the uniform-null expectation.

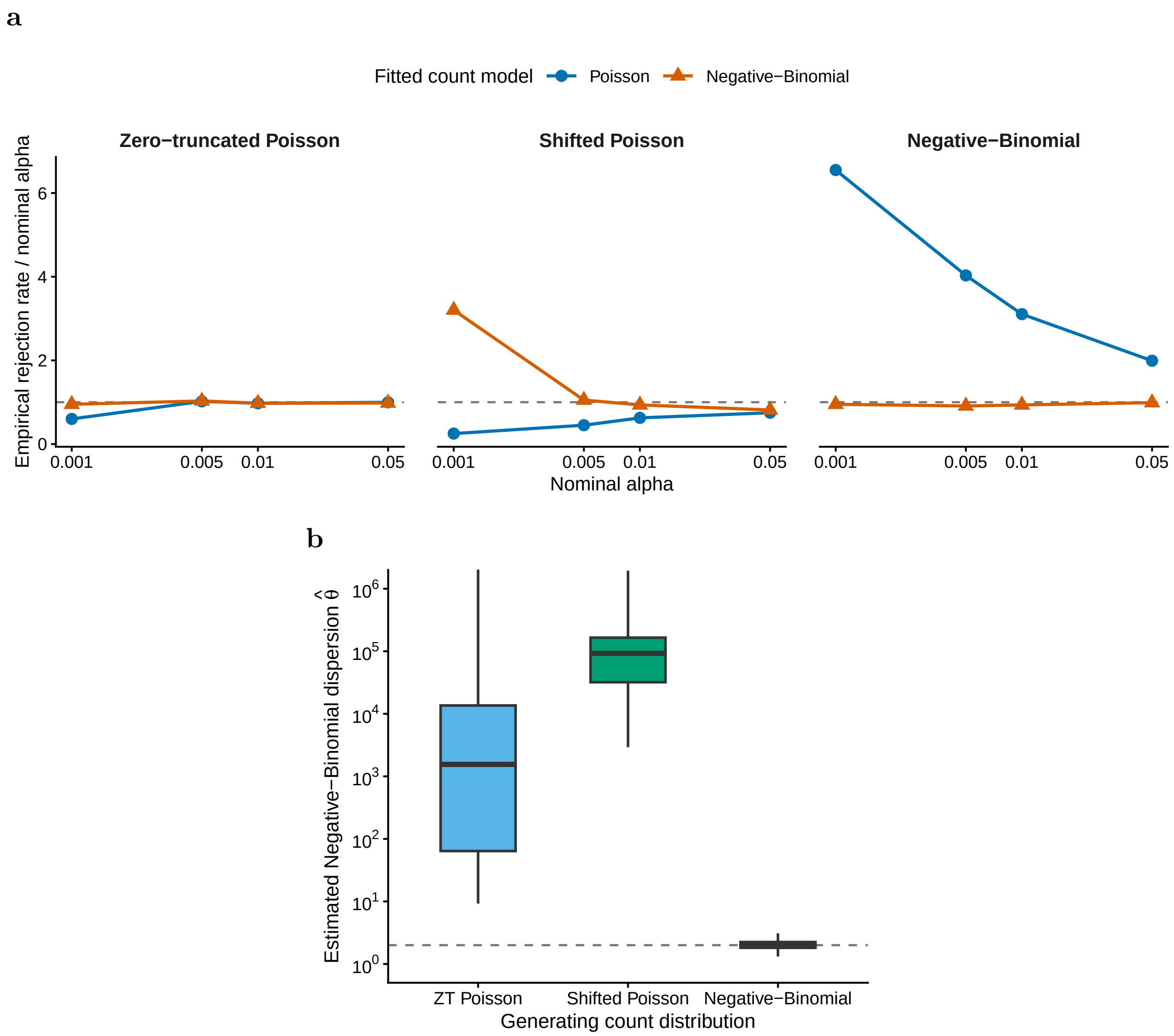


**Supplementary Figure S5:** Count-model calibration in controlled null simulations. (a) Empirical rejection rate divided by nominal $\alpha$ for Poisson and Negative-Binomial Wald tests under the three simulated positive-count distributions. The horizontal dashed line denotes exact calibration. (b) Estimated Negative-Binomial dispersion in the same simulations. Boxplots show medians and interquartile ranges, with whiskers extending to 1.5 times the interquartile range; outliers are omitted and the displayed axis is truncated at $10^6$. The dashed line marks the generating value $\theta = 2$ in the overdispersed regime.

### Joint score test sensitivity analysis

We repeated the pairwise screen in the 10 primary cell types using the two-degree-of-freedom joint score test described above, with all other analysis settings unchanged. For each ordered response-predictor direction, the count and detection score statistics were independent under the factorized hurdle likelihood, and their sum followed a $\chi^2_2$ distribution:

$$T_{\text{joint}} = T_{\text{count}} + T_{\text{detection}} \sim \chi^2_2.$$

Let $M = \sum_c \binom{G_c}{2}$ denote the number of unordered within-chromosome autosomal gene pairs. Because both response-predictor directions were tested, the Bonferroni threshold for each directional test was $0.05/(2M)$. Pairs separated by at most 49 positions in genomic order were evaluated, although the multiple-testing denominator was based on the complete set of $2M$ directional tests. Clusters were called using the same 50-to-2-gene windows and 70% within-window pair-density criterion as in the primary analysis.

The primary and joint-score screens identified 64,295 and 121,821 significant unordered gene pairs, respectively. Of the primary pairs, 58,851 (91.5%) were also identified by the joint-score screen. The two analyses produced 2,485 and 3,036 clusters, respectively. Among the primary clusters, 78.7% had an exact gene-set match in the joint-score analysis. In every cell type, the median maximum Jaccard similarity between a primary cluster and its best joint-score match was 1.0. Furthermore, all four representative cluster gene sets presented in the locus analyses were also recovered by the joint-score approach (Supplementary Table S2). These results indicate that the primary cluster structure was largely preserved, while the joint score test identified additional significant pairs and clusters.

**Supplementary Table S2:** Joint-score sensitivity of pairwise associations and cluster calls. The primary component-union and two-degree-of-freedom joint-score screens were compared across 10 cell types. **a**, Overlap of significant gene pairs; percentages use the corresponding screen as the denominator, and Jaccard is the intersection divided by the union. **b**, Clusters with identical constituent gene sets. **c**, Best Jaccard match to a cluster from the other screen. The **All** row aggregates cell types.

**a. Significant-pair overlap**

| Cell type | Primary pairs | Joint-score pairs | Shared pairs | Primary recovered (%) | Joint shared (%) | Pair Jaccard |
|---|---|---|---|---|---|---|
| B_IN | 3,377 | 6,082 | 3,032 | 89.8 | 49.9 | 0.472 |
| B_Mem | 3,037 | 5,628 | 2,760 | 90.9 | 49.0 | 0.467 |
| CD4_ET | 3,804 | 7,104 | 3,385 | 89.0 | 47.6 | 0.450 |
| CD4_NC | 16,994 | 32,808 | 15,806 | 93.0 | 48.2 | 0.465 |
| CD8_ET | 8,821 | 16,483 | 7,939 | 90.0 | 48.2 | 0.457 |
| CD8_NC | 9,402 | 18,421 | 8,749 | 93.1 | 47.5 | 0.459 |
| CD8_S100B | 2,031 | 3,719 | 1,795 | 88.4 | 48.3 | 0.454 |
| Mono_C | 5,210 | 9,765 | 4,789 | 91.9 | 49.0 | 0.470 |
| Mono_NC | 1,846 | 3,345 | 1,594 | 86.3 | 47.7 | 0.443 |
| NK | 9,773 | 18,466 | 9,002 | 92.1 | 48.7 | 0.468 |
| **All** | **64,295** | **121,821** | **58,851** | **91.5** | **48.3** | **0.462** |

**b. Exact cluster overlap**

| Cell type | Primary clusters | Joint-score clusters | Exact clusters | Primary exact (%) | Joint exact (%) |
|---|---|---|---|---|---|
| B_IN | 172 | 214 | 140 | 81.4 | 65.4 |
| B_Mem | 171 | 215 | 146 | 85.4 | 67.9 |
| CD4_ET | 201 | 232 | 168 | 83.6 | 72.4 |
| CD4_NC | 513 | 631 | 369 | 71.9 | 58.5 |
| CD8_ET | 316 | 406 | 243 | 76.9 | 59.9 |
| CD8_NC | 360 | 438 | 290 | 80.6 | 66.2 |
| CD8_S100B | 123 | 147 | 108 | 87.8 | 73.5 |
| Mono_C | 188 | 223 | 146 | 77.7 | 65.5 |
| Mono_NC | 101 | 124 | 85 | 84.2 | 68.5 |
| NK | 340 | 406 | 261 | 76.8 | 64.3 |
| **All** | **2,485** | **3,036** | **1,956** | **78.7** | **64.4** |

**c. Best-match cluster overlap**

| Cell type | Primary median best Jaccard | Primary best Jaccard $\geq 0.5$ (%) | Joint median best Jaccard | Joint best Jaccard $\geq 0.5$ (%) |
|---|---|---|---|---|
| B_IN | 1.000 | 86.6 | 1.000 | 69.6 |
| B_Mem | 1.000 | 88.3 | 1.000 | 70.2 |
| CD4_ET | 1.000 | 87.1 | 1.000 | 75.0 |
| CD4_NC | 1.000 | 80.7 | 1.000 | 65.6 |
| CD8_ET | 1.000 | 84.2 | 1.000 | 65.0 |
| CD8_NC | 1.000 | 86.9 | 1.000 | 70.8 |
| CD8_S100B | 1.000 | 91.1 | 1.000 | 76.2 |
| Mono_C | 1.000 | 83.0 | 1.000 | 70.0 |
| Mono_NC | 1.000 | 88.1 | 1.000 | 71.8 |
| NK | 1.000 | 82.4 | 1.000 | 69.0 |
| **All** | **1.000** | **84.7** | **1.000** | **69.1** |

### Cluster annotation enrichment

To assess whether sc-pcQTL clusters captured shared functional or regulatory features, we compared clusters containing 2–5 genes with neighboring null gene sets sampled to match the observed size distribution within each primary cell type. Clusters containing more than five genes were excluded from this enrichment analysis but were retained in all other analyses.

For each binary annotation, we fitted the following logistic-regression model:

$$\text{annotation status} \sim \text{correlated cluster} + \text{number of genes} + \log_{10}(\text{cluster length}),$$

where cluster length was defined as the genomic span of the gene set. We reported the odds ratio for correlated-cluster status and adjusted the corresponding $p$-values across annotation and correlation-type strata. Annotation categories were excluded when any expected frequency in the corresponding contingency table was below 1. Tested features included shared functional annotations, such as paralogy and Gene Ontology biological-process terms; shared ABC enhancer-gene links; promoter-orientation and gene-overlap configurations; and whether gene sets spanned CTCF peaks or topologically associating domain boundaries.

Gene coordinates and strand came from GENCODE v19. Gene Ontology biological-process terms and human paralogs came from the GO Annotation database and Ensembl BioMart, respectively. ABC enhancer-gene links came from Nasser et al., whereas GM12878 ENCODE CTCF peaks and Hi-C TADs provided boundary annotations.

In the analysis pooling all correlation types, sc-pcQTL clusters were enriched for paralogous genes, shared Gene Ontology biological-process terms, same-strand promoter configurations, and same-strand gene overlaps. They were also less likely than matched neighboring null sets to span CTCF peaks or topologically associating domain boundaries.

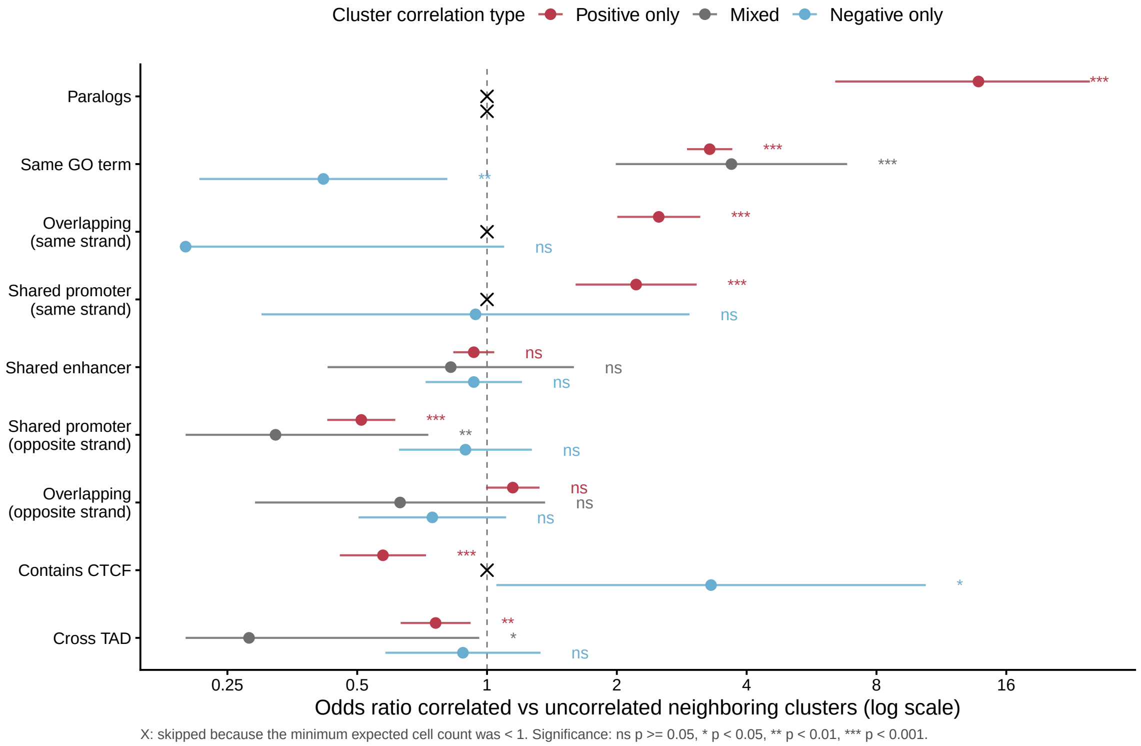


**Supplementary Figure S6:** Size- and length-adjusted annotation enrichment of correlated sc-pcQTL clusters in the 10 primary cell types, stratified by cluster correlation type. Logistic-regression odds ratios (log scale) with 95% confidence intervals compare 2,464 enrichment-eligible clusters of 2–5 genes against 87,714 neighboring null gene sets sampled to match the correlated clusters' size distribution. Models adjust for gene number and cluster length; crosses mark tests skipped for expected frequency below one. In the all-cluster analysis, correlated neighboring-gene clusters were enriched for shared Gene Ontology biological-process terms (OR = 2.81, 95% CI 2.51–3.16), paralogs (OR = 11.2), same-strand promoter architecture (OR = 1.89), and same-strand overlap (OR = 2.20). They were depleted for crossing CTCF peaks (OR = 0.67) and TAD boundaries (OR = 0.76).

### Cross-mappability sensitivity analysis

To assess whether ambiguous read mapping affected the detected co-expression clusters or pcQTL signals, we used the hg19/GENCODE v19 symmetric-mean cross-mappability resource (Saha and Battle, 2018). Following the published pcQTL workflow for bulk level data (Lawrence et al., 2026), a gene pair was considered cross-mappable when its score exceeded 100, and a cluster was considered cross-mappable if it contained at least one such pair. We tested whether cross-mappable pairs were enriched in sc-pcQTL clusters, compared the proportion of cluster-PC phenotypes yielding a SuSiE credible set between clusters with and without cross-mappable pairs, repeated the pcQTL and colocalization summaries after excluding cross-mappable clusters, and examined all within-cluster pairs at the *GIMAP* locus.

A minority of clusters met the cross-mappability criterion. At the primary threshold (> 100), cross-mappable gene pairs were enriched in sc-pcQTL clusters (odds ratio = 4.01, 95% CI 3.51–4.59). Excluding cross-mappable clusters retained 87.2% of significant pcQTL phenotypes and 89.1% of pcQTL-specific colocalization results while preserving the relative pcQTL-specific increment (Supplementary Figure S7). All 21 gene pairs in the *GIMAP* cluster had a cross-mappability score of zero (Supplementary Table S3).

**Supplementary Table S3:** Cross-mappability audit of the representative *GIMAP* cluster. All seven genes were uniquely matched to GENCODE v19, making all 21 internal gene pairs evaluable. The hg19 symmetric-mean resource was computed from exon 75-mers and UTR 36-mers while allowing up to two mismatches and contains only non-zero pairs. None of the 21 pairs was present in the sparse table, so each was assigned a score of zero following the published pcQTL workflow; no pair exceeded the primary threshold of 100.

| Gene pair | Score | Gene pair | Score |
|---|---|---|---|
| *GIMAP8–GIMAP7* | 0 | *GIMAP4–GIMAP6* | 0 |
| *GIMAP8–GIMAP4* | 0 | *GIMAP4–GIMAP2* | 0 |
| *GIMAP8–GIMAP6* | 0 | *GIMAP4–GIMAP1* | 0 |
| *GIMAP8–GIMAP2* | 0 | *GIMAP4–GIMAP5* | 0 |
| *GIMAP8–GIMAP1* | 0 | *GIMAP6–GIMAP2* | 0 |
| *GIMAP8–GIMAP5* | 0 | *GIMAP6–GIMAP1* | 0 |
| *GIMAP7–GIMAP4* | 0 | *GIMAP6–GIMAP5* | 0 |
| *GIMAP7–GIMAP6* | 0 | *GIMAP2–GIMAP1* | 0 |
| *GIMAP7–GIMAP2* | 0 | *GIMAP2–GIMAP5* | 0 |
| *GIMAP7–GIMAP1* | 0 | *GIMAP1–GIMAP5* | 0 |
| *GIMAP7–GIMAP5* | 0 | | |

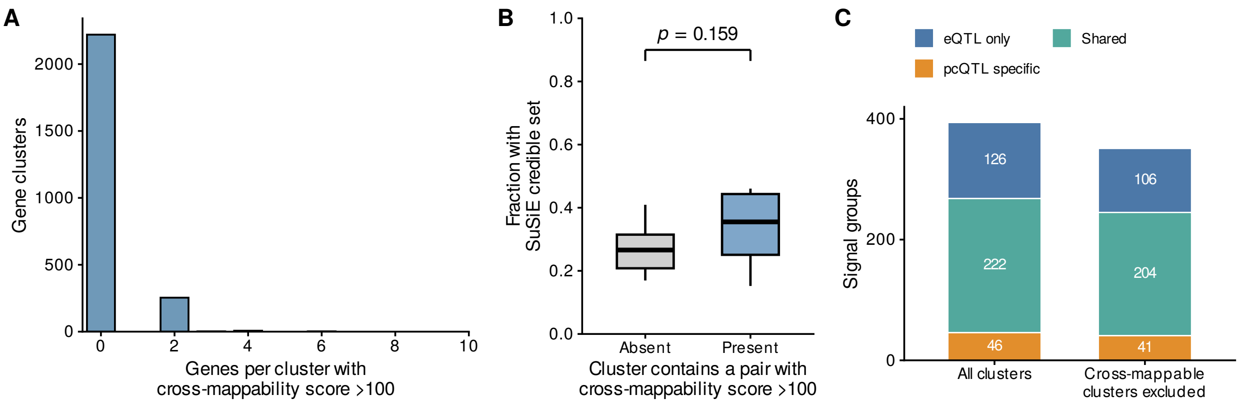


**Supplementary Figure S7:** Cross-mappability sensitivity analysis in the 10 primary cell types. (A) Number of genes per cluster involved in at least one gene pair with a symmetric-mean cross-mappability score greater than 100. (B) Fraction of cluster-PC phenotypes with a retained SuSiE credible set in clusters with or without at least one pair exceeding this threshold. Boxes show the median and interquartile range, and the displayed $p$-value is from a two-sided Welch $t$-test across cell types. (C) Strict colocalization classes before and after excluding cross-mappable clusters.

### ACAT sensitivity analysis

To obtain a single phenotype-level association test, we combined the *cis*-variant $p$-values within each successfully tested cluster-PC phenotype using the aggregated Cauchy association test (ACAT) (Liu et al., 2019). The Cauchy combination is robust to correlation among variant-level tests and follows the gene-level aggregation strategy used in SAIGE-QTL (Zhou et al., 2024). We then applied the Benjamini-Hochberg procedure across phenotypes. ACAT identified 2,017 significant phenotypes, including 1,992 of the 2,040 primary discoveries (97.6%; Supplementary Table S4), demonstrating strong concordance between the primary variant-level discovery criterion and the ACAT-based phenotype-level analysis.

**Supplementary Table S4:** Concordance of primary and ACAT-BH pcQTL discovery criteria. Primary calls required at least one cis variant with within-phenotype Benjamini–Hochberg $q < 0.05$. For the secondary analysis, the aggregated Cauchy association test (ACAT) combined all cis-variant $p$-values for each cluster-PC phenotype, followed by Benjamini–Hochberg correction across the 4,353 successfully tested phenotypes. Both denotes phenotypes significant by both criteria.

| Cell type | Successfully tested | Primary FDR | ACAT-BH | Both | Primary only | ACAT-BH only |
|---|---|---|---|---|---|---|
| B_IN | 299 | 110 | 108 | 107 | 3 | 1 |
| B_Mem | 302 | 101 | 99 | 98 | 3 | 1 |
| CD4_ET | 341 | 142 | 137 | 136 | 6 | 1 |
| CD4_NC | 957 | 556 | 548 | 544 | 12 | 4 |
| CD8_ET | 562 | 269 | 267 | 265 | 4 | 2 |
| CD8_NC | 632 | 278 | 278 | 274 | 4 | 4 |
| CD8_S100B | 208 | 72 | 73 | 69 | 3 | 4 |
| Mono_C | 299 | 110 | 109 | 107 | 3 | 2 |
| Mono_NC | 166 | 71 | 69 | 67 | 4 | 2 |
| NK | 587 | 331 | 329 | 325 | 6 | 4 |
| **All** | **4,353** | **2,040** | **2,017** | **1,992** | **48** | **25** |

### SuSiE colocalization sensitivity analyses

We used `coloc.susie` to evaluate colocalization between allele-compatible OneK1K QTL signals and GWAS fine-mapped signals in the FinnGen study. Within each cell type and local gene cluster, fine-mapped QTL and GWAS signals connected through qualifying QTL-GWAS or eQTL-pcQTL colocalization links formed colocalized QTL-GWAS signal groups (hereafter referred to as signal groups). Each retained signal group contained at least one GWAS signal and was classified by its QTL composition. At PPH4 thresholds of 0.70, 0.75, and 0.80, the analysis identified 49, 46, and 38 pcQTL-specific signal groups and 377, 348, and 296 eQTL-containing signal groups, respectively. Overall, the pcQTL-specific class was retained across all three thresholds.

As an additional sensitivity analysis, we applied `coloc.abf` (Giambartolomei et al., 2014) to a broader set of observed shared variants under the single-causal-variant assumption (Supplementary Figure S9).

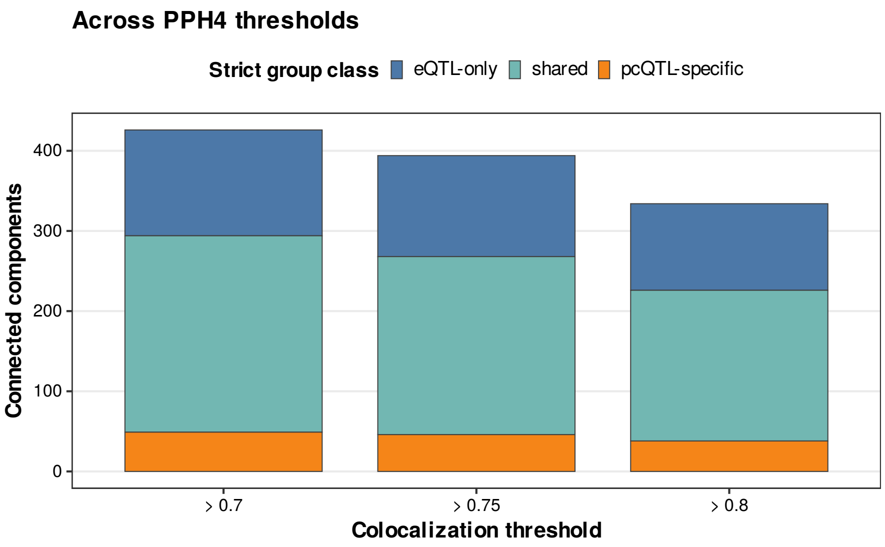


**Supplementary Figure S8:** Classes of signal groups across SuSiE colocalization thresholds. Stacked bars show eQTL-only, shared, and pcQTL-specific signal groups obtained after fine-mapping and `coloc.susie` grouping at PPH4 > 0.70, > 0.75, and > 0.80.

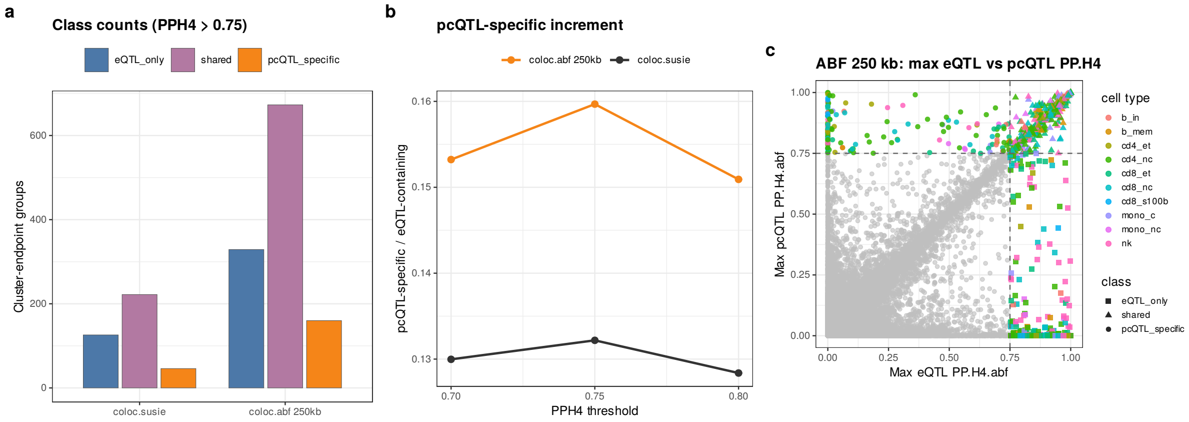


**Supplementary Figure S9:** Observed-shared-variant `coloc.abf` sensitivity compared with `coloc.susie` for identifying signal groups in the 10 primary cell types. (a) Class counts at PPH4 > 0.75 for the SuSiE signal-group analysis and the 250 kb QTL-lead-centered ABF analysis. (b) Relative pcQTL-specific increment, defined as pcQTL-specific divided by eQTL-containing groups, across posterior thresholds. (c) Maximum eQTL versus pcQTL PPH4 for 250 kb ABF cluster-trait groups; grey points are non-colocalized groups and dashed lines mark PPH4 = 0.75. The ABF analysis uses a broader candidate set and assumes at most one causal variant per trait in each region, so it provides sensitivity context for the primary fine-mapped analysis.

### PIP-weighted nominal effects and regulatory annotations

Signal groups were linked to their corresponding QTL credible sets. For each credible set CS and constituent gene $g$ with available nominal eQTL statistics, $\mathcal{M}_{g,\mathrm{CS}}$ denotes the credible-set variants matched by chromosome, position, and alleles. Nominal effect estimates were aligned to the QTL effect allele, with the sign reversed for swapped alleles. We defined the PIP-weighted nominal gene effect as

$$\hat{\beta}_{g,\mathrm{CS}} = \frac{\sum_{j\in\mathcal{M}_{g,\mathrm{CS}}} \mathrm{PIP}_j \hat{\beta}^*_{jg}}{\sum_{j\in\mathcal{M}_{g,\mathrm{CS}}} \mathrm{PIP}_j},$$

where $\mathrm{PIP}_j$ is the marginal SuSiE posterior inclusion probability of variant $j$, and $\hat{\beta}^*_{jg}$ is its allele-aligned nominal effect on gene $g$. Each credible set was summarized by

$$\max_g |\hat{\beta}_{g,\mathrm{CS}}| \qquad \text{and} \qquad \mathrm{CV}_{\mathrm{CS}} = \frac{\mathrm{SD}_g(|\hat{\beta}_{g,\mathrm{CS}}|)}{\mathrm{Mean}_g(|\hat{\beta}_{g,\mathrm{CS}}|)},$$

where the standard deviation and mean were calculated across constituent genes with finite effect estimates. These quantities are nominal expression-effect summaries rather than calibrated allelic fold-change estimates.

For regulatory annotation $a$, we calculated

$$A_{a,\mathrm{CS}} = \frac{\sum_{j\in\mathrm{CS}} \mathrm{PIP}_j I_{ja}}{\sum_{j\in\mathrm{CS}} \mathrm{PIP}_j},$$

where $I_{ja} = 1$ when variant $j$ overlaps annotation $a$ and 0 otherwise.

pcQTL-specific credible sets had lower maximum absolute PIP-weighted nominal effects and lower cross-gene CV than non-colocalized and eQTL-only credible sets (Supplementary Figure S10).

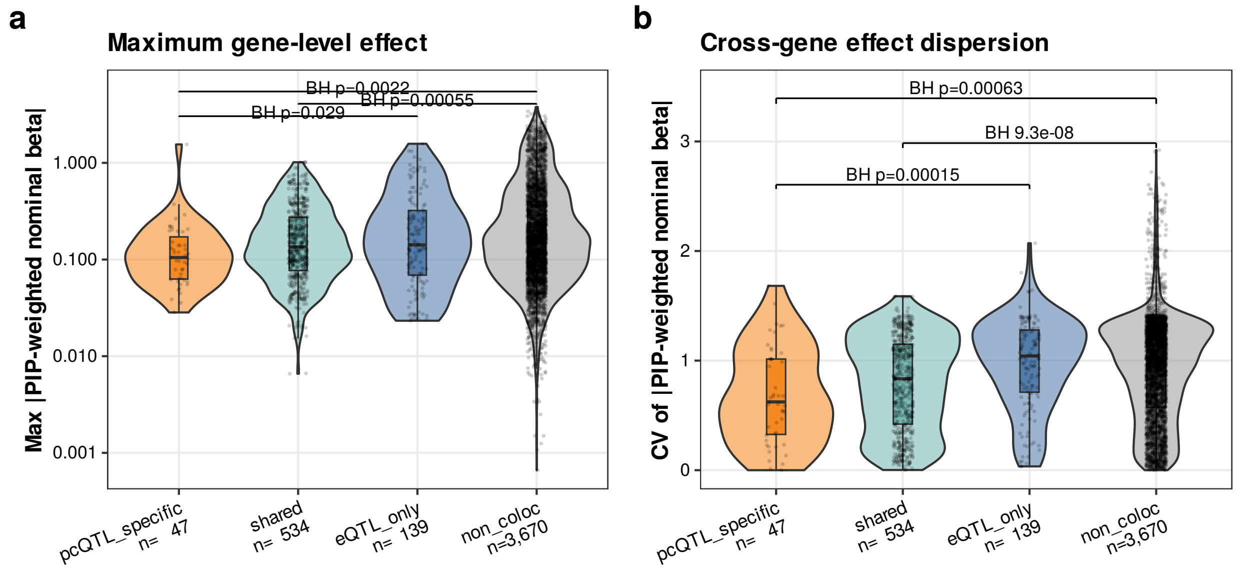


**Supplementary Figure S10:** PIP-weighted nominal gene-effect profiles across signal-group classes and the non-colocalized reference class in the 10 primary cell types. (a) Maximum absolute PIP-weighted nominal effect by class. (b) Cross-gene nominal-effect coefficient of variation by class. Violin/box plots show finite QTL credible sets (pcQTL-specific n=47, shared n=534, eQTL-only n=139, non-colocalized n=3,670). Brackets report Benjamini-Hochberg-adjusted pairwise Wilcoxon $p$-values.

### PC-loading concentration among pcQTL-specific signal groups

The top squared PC-loading share was $\max_g(l_g^2)/\sum_g l_g^2$. A pcQTL-specific signal group containing multiple pcQTL signals was classified as meeting the 75% or 90% loading-share threshold only when every contributing cluster-PC phenotype met that threshold. For each credible set, the largest-gene effect fraction was $\max_g(|\hat{\beta}_{g,\mathrm{CS}}|)/\sum_g |\hat{\beta}_{g,\mathrm{CS}}|$.

**Supplementary Table S5:** PC-loading and PIP-weighted nominal-effect concentration among pcQTL-specific signal groups.

| Summary measure | Analysis unit | Result |
|---|---|---|
| Total pcQTL-specific signal groups | Signal group | 46 |
| Signal groups arising from two-gene clusters | Signal group | 38/46 (82.6%) |
| Top squared PC-loading share $\geq$ 75% | Signal group | 45/46 (97.8%) |
| Top squared PC-loading share $\geq$ 90% | Signal group | 40/46 (87.0%) |
| Median top squared PC-loading share | Signal group | 99.9% |
| Median largest-gene fraction of the absolute PIP-weighted nominal effect | Credible set | 70.7% ($n = 47$) |

### S-LDSC heritability enrichment and trait-level context

We used stratified LD score regression (S-LDSC) (Bulik-Sullivan et al., 2015; Finucane et al., 2015; Gazal et al., 2017) to compare disease-heritability enrichment between pcQTL and single-gene eQTL annotations across 247 immune-related traits in the FinnGen study.

For regression SNP $j$, S-LDSC models the expected GWAS association statistic as

$$\mathrm{E}[\chi_j^2] = 1 + Na + N \sum_C \tau_C \ell(j, C), \qquad \ell(j, C) = \sum_k A_C(k) r_{jk}^2,$$

where $N$ is the GWAS sample size, $a$ captures confounding, $\tau_C$ is the conditional effect of annotation $C$, and $\ell(j, C)$ is its LD score. Binary annotations pooled FDR-significant *cis*-QTL variants (MAF $\geq 0.05$) across the 10 primary cell types and were matched to the 1000 Genomes Phase 3 European reference (The 1000 Genomes Project Consortium, 2015). Models were fitted using baseline-LD v2.2 after removing its molecular-QTL MaxCPP annotations (Hormozdiari et al., 2018), and the extended MHC was excluded. Heritability quality control and genetic-correlation clumping at $|r_g| \geq 0.7$ retained 35 approximately independent traits.

For a binary annotation $C$, heritability enrichment was defined as

$$\mathrm{Enrichment}_C = \frac{h_g^2(C)/h_g^2}{M_C/M},$$

where $h_g^2(C)/h_g^2$ and $M_C/M$ denote the proportions of heritability and reference variants attributable to the annotation, respectively. The standardized effect was

$$\tau_C^* = \frac{M \,\mathrm{sd}(C)\, \tau_C}{h_g^2},$$

(Gazal et al., 2017). Marginal and joint annotation models were combined across traits by random-effects meta-analysis (DerSimonian and Laird, 1986; Hujoel et al., 2019).

pcQTL and eQTL annotations showed 1.80× and 1.64× heritability enrichment, respectively. In marginal models that included one QTL annotation at a time, the baseline-LD-adjusted pcQTL effect was not significant ($\tau^* = 0.020$, $p = 0.296$), whereas the eQTL effect was positive ($\tau^* = 0.065$, $p = 3.38 \times 10^{-4}$). In the joint model, the pcQTL effect remained close to zero ($\tau^* = 0.004$, $p = 0.835$), whereas the eQTL effect remained positive ($\tau^* = 0.046$, $p = 0.0137$; Supplementary Figure S11; Supplementary Table S6).

**Supplementary Table S6:** Primary S-LDSC meta-analysis across 35 approximately independent disease traits. Marginal models included baseline-LD and one QTL annotation at a time and report heritability enrichment and standardized effects ($\tau^*$); the joint model included both QTL annotations. $\Delta\tau^*$ is the pcQTL-minus-eQTL contrast. Confidence intervals are 95%. The complete per-trait and secondary-analysis results are provided in the machine-readable S-LDSC result files archived with the Supplementary Data.

| Model | Annotation or contrast | Metric | Estimate (95% CI) | $p$-value |
|---|---|---|---|---|
| Marginal | pcQTL | Enrichment | 1.804 (1.524, 2.085) | — |
| Marginal | eQTL | Enrichment | 1.642 (1.525, 1.759) | — |
| Marginal | pcQTL | $\tau^*$ | 0.020 (−0.017, 0.057) | 0.296 |
| Marginal | eQTL | $\tau^*$ | 0.065 (0.030, 0.101) | $3.38 \times 10^{-4}$ |
| Joint | pcQTL | $\tau^*$ | 0.004 (−0.035, 0.043) | 0.835 |
| Joint | eQTL | $\tau^*$ | 0.046 (0.009, 0.082) | 0.0137 |
| Joint contrast | pcQTL minus eQTL | $\Delta\tau^*$ | −0.029 (−0.093, 0.034) | 0.362 |

Enrichment is summarized by its confidence interval; $p$-values are shown for standardized effects and their contrast.

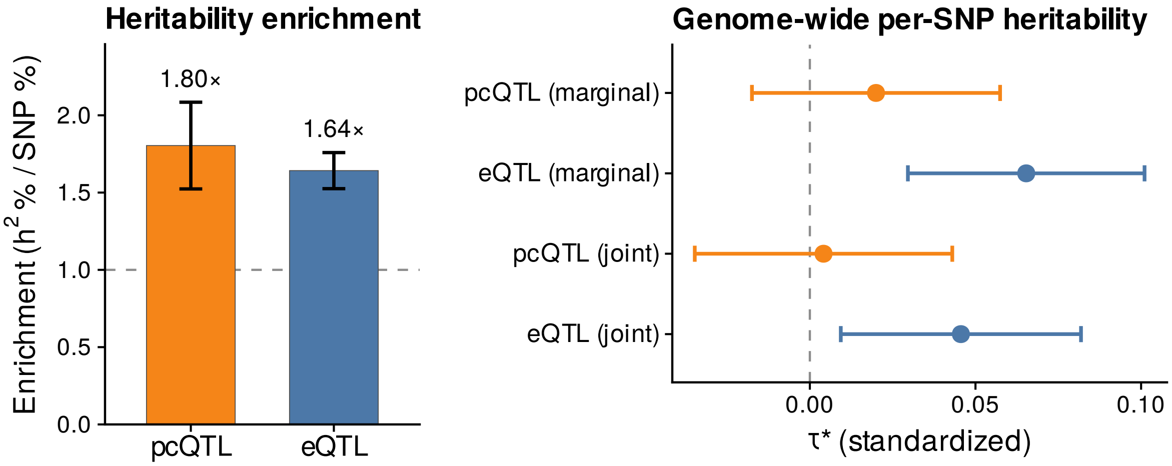


**Supplementary Figure S11:** Disease heritability enrichment of pcQTL and single-gene eQTL annotations. The primary S-LDSC meta-analysis included 35 approximately independent disease traits in the FinnGen study selected from 247 pre-specified traits after SNP-heritability quality control and genetic-correlation clumping. Annotations comprise all per-feature FDR-significant *cis*-QTL variants with MAF $\geq 0.05$, pooled across the 10 primary cell types. Models were conditioned on baseline-LD v2.2 after removal of its built-in molecular-QTL annotations. Error bars show 95% confidence intervals. Left, marginal heritability enrichment ($h^2\%/\text{SNP}\%$); right, marginal and jointly conditional standardized per-SNP effects, $\tau^*$. The joint pcQTL-minus-eQTL difference was $\Delta\tau^* = -0.029$ (95% CI, $-0.093$ to 0.034; $p = 0.362$).

### Follow-up analyses of the representative *GIMAP* locus

We performed three complementary follow-up analyses to further characterize the *GIMAP* signal. First, we compared recovery of the seven-gene cluster and pcQTL colocalization across the 10 primary cell types to assess cell-type specificity. Second, we constructed donor-level pseudobulk mixtures containing increasing proportions of $CD8^+$ naïve and central-memory T cells (CD8_NC) to evaluate the effect of cell-type mixing. Third, we performed SMR analyses to assess gene-level evidence for both the cluster-PC phenotype and the lymphocyte-count GWAS signal.

The complete seven-gene *GIMAP* cluster was recovered in several lymphoid cell types, indicating that the local co-expression structure was not unique to CD8_NC cells. However, colocalization with the lymphocyte-count GWAS signal was observed only for the pcQTL in CD8_NC cells (PPH4 = 0.93); the next-highest value was PPH4 = 0.66 in CD4_ET cells. Thus, recovery of the same co-expression module across cell types did not necessarily imply colocalization of the corresponding cluster-PC phenotype with lymphocyte count.

For each pseudobulk mixture, we remapped cluster-PC pcQTLs and constituent-gene eQTLs, fine-mapped them using OneK1K LD, and tested colocalization with the same lymphocyte-count GWAS signal in the FinnGen study used in the primary analysis. The pcQTL-GWAS comparison exceeded the primary threshold only in the mixture composed of 100% CD8_NC cells (PPH4 = 0.82). Under the observed cell-type composition and in mixtures containing up to 75% CD8_NC cells, PPH4 remained at or below 0.16, whereas all constituent-gene eQTLs remained below the threshold. This pattern is consistent with attenuation of the cluster-PC colocalization signal by cell-type mixing.

In SMR analyses of the cluster-PC phenotype, five constituent genes met the Bonferroni-corrected significance threshold and showed no evidence of heterogeneity in the HEIDI test. In the gene-level analysis of lymphocyte count, *GIMAP7* showed the strongest HEIDI-consistent association ($p_{\mathrm{SMR}} = 6.41 \times 10^{-4}$, $p_{\mathrm{HEIDI}} = 0.217$), whereas the associations for *GIMAP4* and *GIMAP2* showed evidence of HEIDI heterogeneity (Supplementary Figure S12d,e). These SMR and HEIDI results provide descriptive gene-level follow-up but do not identify a single causal mediator. Together, the cross-cell-type, pseudobulk-mixture, and gene-level analyses support a cell-type-specific, multi-gene interpretation of the *GIMAP* signal.

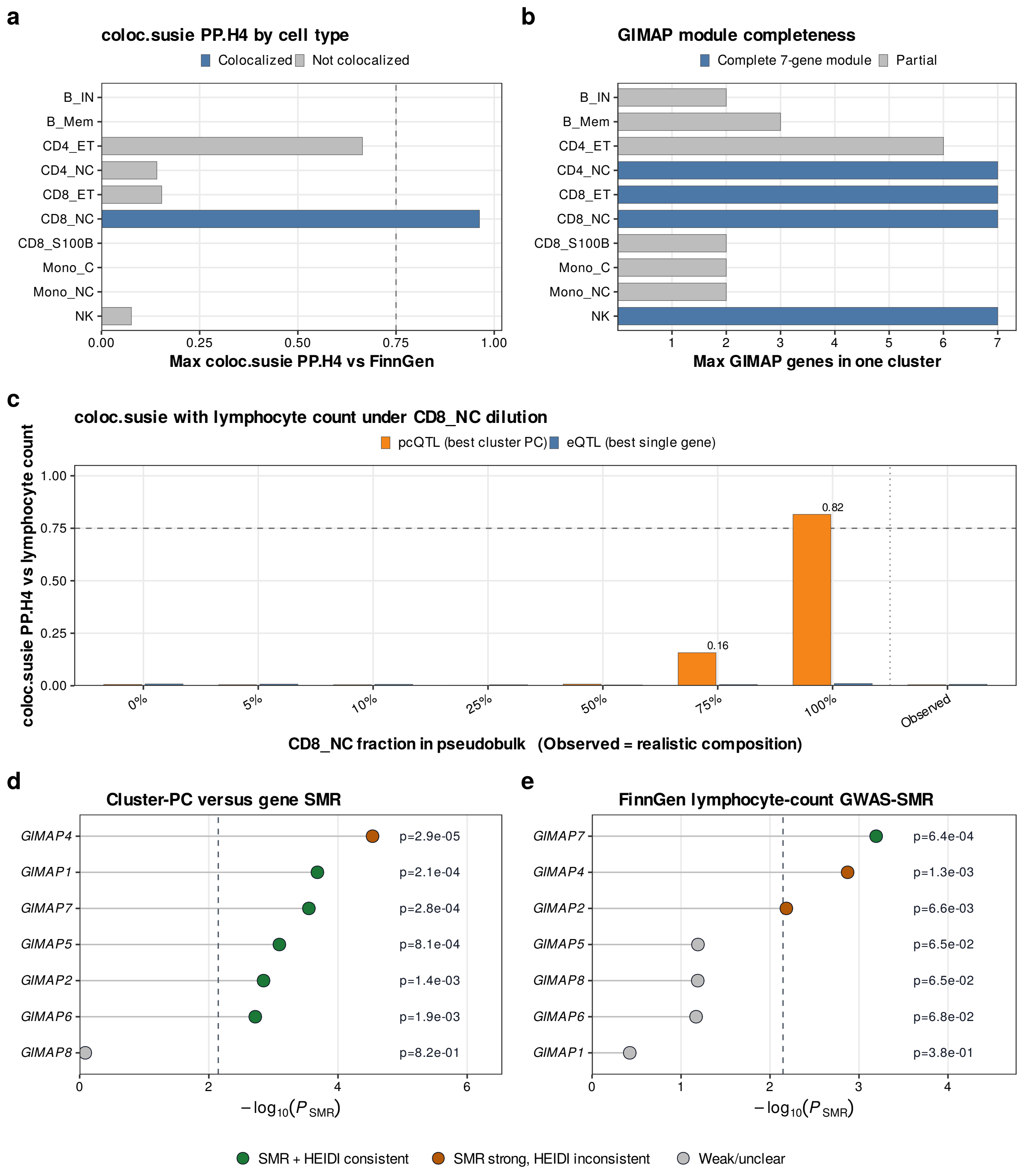


**Supplementary Figure S12:** Representative *GIMAP* locus follow-up. (a) Maximum `coloc.susie` PPH4 for the *GIMAP* cluster-PC pcQTL across cell types; only the pcQTL in CD8_NC cells exceeded the PPH4 = 0.75 threshold. (b) Maximum number of *GIMAP* genes recovered within a single local cluster in each cell type. (c) PPH4 for the best cluster-PC pcQTL and best same-cluster single-gene eQTL across donor-level pseudobulk mixtures containing 0–100% CD8_NC cells and the observed cell-type composition. Each mixture QTL was fine-mapped using OneK1K LD and compared with the same lymphocyte-count GWAS signal in the FinnGen study used in the primary analysis. The dashed line marks PPH4 = 0.75. (d) SMR results for the cluster-PC phenotype and constituent genes. (e) SMR results for the lymphocyte-count GWAS in the FinnGen study. Dashed vertical lines indicate the seven-gene Bonferroni threshold. Colors distinguish Bonferroni-significant associations with or without HEIDI consistency and associations that did not pass the significance threshold. HEIDI consistency was defined as $p_{\mathrm{HEIDI}} > 0.01$ using at least three variants. SMR and HEIDI were used as descriptive gene-level follow-up for this multi-gene locus.

### Additional representative loci

Three additional pcQTL-specific loci satisfied the primary SuSiE colocalization and shared-posterior-mass criteria. Regional plots are shown using hg38 coordinates, the full OneK1K fine-mapping window, and a common 1000 Genomes European LD reference for visualization.

The *ASCL2-C11orf21-TSPAN32* locus is supported by T-cell biology, the *EIF3K-ACTN4* locus is linked to platelet cytoskeletal biology, and the *EVI2B-EVI2A* locus has established hematopoietic relevance (Liu et al., 2014; Lombardo et al., 2019; O'Sullivan et al., 2021; Zjablovskaja et al., 2017).

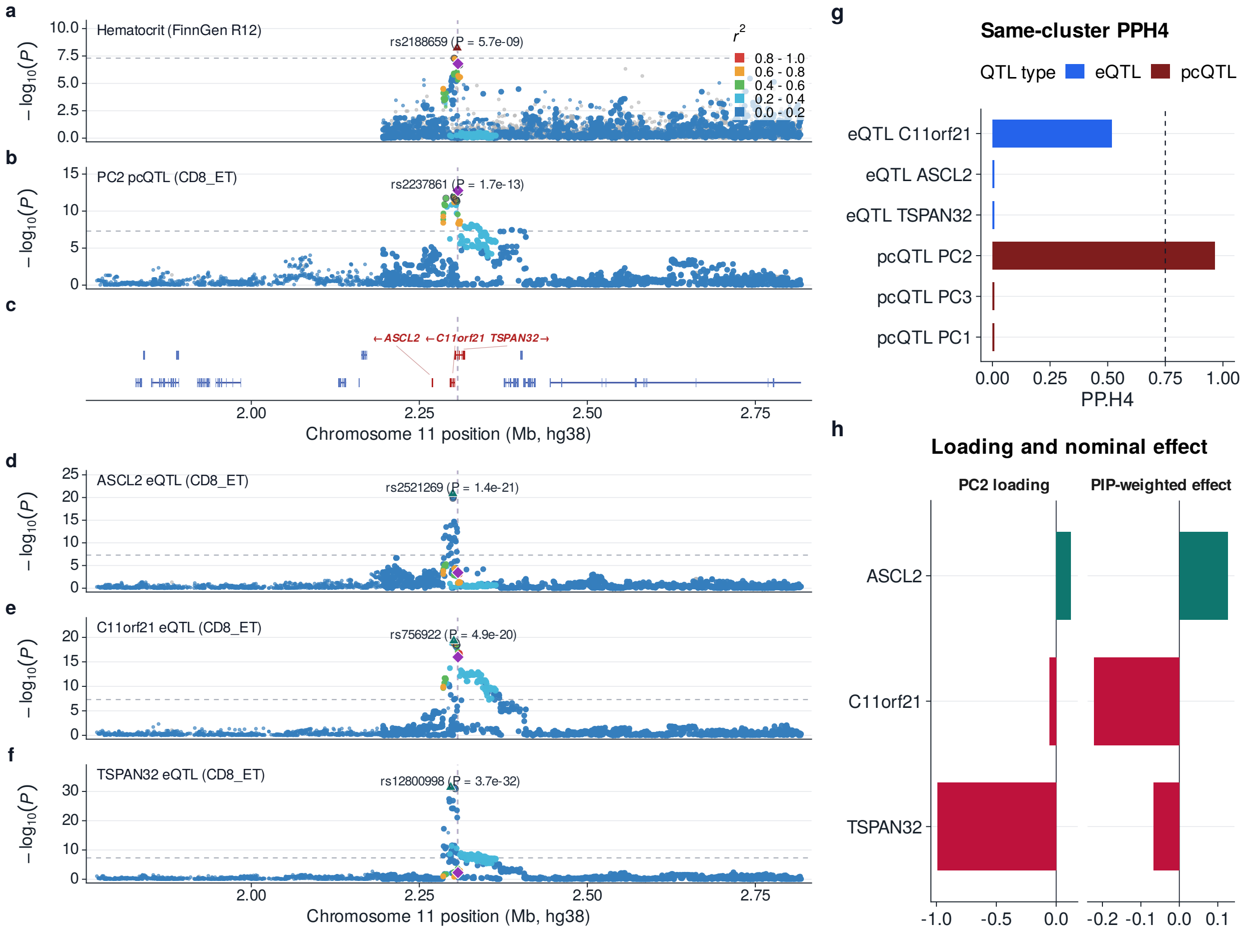


**Supplementary Figure S13:** Representative locus view for CD8$^+$ T cells with an effector-memory phenotype (CD8_ET), showing the *ASCL2-C11orf21-TSPAN32* PC2 pcQTL and hematocrit GWAS in the FinnGen study. The pcQTL-GWAS comparison passed the main threshold (PPH4 = 0.964, PPH3 = 0.036) and shared-posterior-mass QC across 902 allele-compatible shared variants. Panels show regional GWAS in the FinnGen, OneK1K pcQTL, gene context, all same-cluster single-gene eQTL regional associations, same-cluster QTL-GWAS PPH4, and PIP-weighted nominal-effect summaries. LD colors are a display-only layer from a unified 1000 Genomes Phase 3 EUR reference.

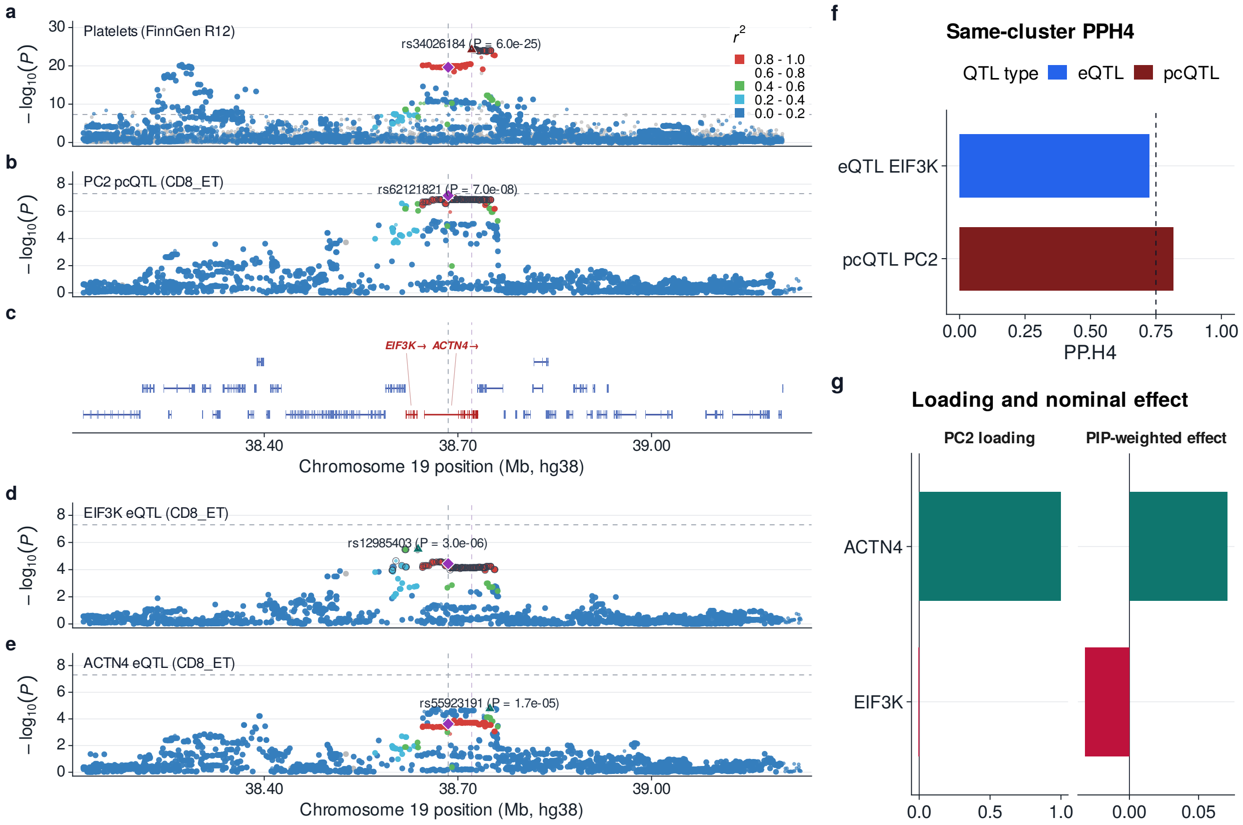


**Supplementary Figure S14:** Representative locus view for CD8$^+$ T cells with an effector-memory phenotype (CD8_ET), showing the *EIF3K-ACTN4* PC2 pcQTL and platelet-count GWAS in the FinnGen study. The pcQTL-GWAS comparison passed the main threshold (PPH4 = 0.817, PPH3 = 0.182) and shared-posterior-mass QC across 1,633 allele-compatible shared variants. Panels show regional GWAS in the FinnGen study, OneK1K pcQTL, gene context, all same-cluster single-gene eQTL regional associations, same-cluster QTL-GWAS PPH4, and PIP-weighted nominal-effect summaries. LD colors are a display-only layer from a unified 1000 Genomes Phase 3 EUR reference.

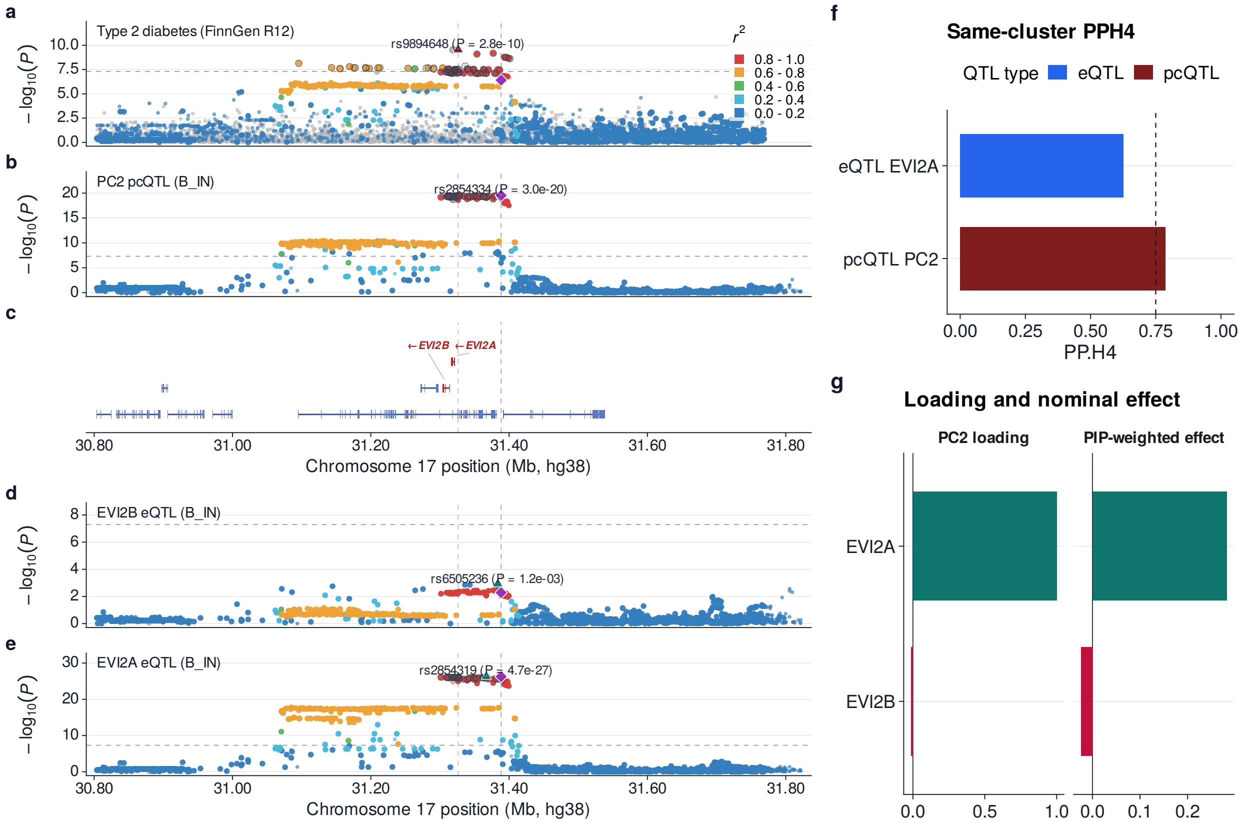


**Supplementary Figure S15:** Representative locus view for immature and naïve B cells (B_IN), showing the *EVI2B*-*EVI2A* PC2 pcQTL and type 2 diabetes GWAS in the FinnGen study. The pcQTL-GWAS comparison passed the main threshold (PPH4 = 0.787, PPH3 = 0.213) and shared-posterior-mass QC across 905 allele-compatible shared variants. Panels show regional GWAS in the FinnGen study, OneK1K pcQTL, gene context, all same-cluster single-gene eQTL regional associations, same-cluster QTL-GWAS PPH4, and PIP-weighted nominal-effect summaries. LD colors are a display-only layer from a unified 1000 Genomes Phase 3 EUR reference.

### References


B. K. Bulik-Sullivan et al. LD Score regression distinguishes confounding from polygenicity in genome-wide association studies. *Nature Genetics*, 47(3):291–295, 2015. doi: 10.1038/ng.3211.

R. DerSimonian and N. Laird. Meta-analysis in clinical trials. *Controlled Clinical Trials*, 7(3): 177–188, 1986. doi: 10.1016/0197-2456(86)90046-2.

H. K. Finucane et al. Partitioning heritability by functional annotation using genome-wide association summary statistics. *Nature Genetics*, 47(11):1228–1235, 2015. doi: 10.1038/ng.3404.

S. Gazal et al. Linkage disequilibrium-dependent architecture of human complex traits shows action of negative selection. *Nature Genetics*, 49(10):1421–1427, 2017. doi: 10.1038/ng.3954.

C. Giambartolomei et al. Bayesian test for colocalisation between pairs of genetic association studies using summary statistics. *PLOS Genetics*, 10(5):e1004383, 2014. doi: 10.1371/journal.pgen.10043 83.

F. Hormozdiari et al. Leveraging molecular quantitative trait loci to understand the genetic architecture of diseases and complex traits. *Nature Genetics*, 50(7):1041–1047, 2018. doi: 10.1038/s41588-018-0148-2.

M. L. A. Hujoel, S. Gazal, F. Hormozdiari, B. van de Geijn, and A. L. Price. Disease heritability enrichment of regulatory elements is concentrated in elements with ancient sequence age and conserved function across species. *The American Journal of Human Genetics*, 104(4):611–624, 2019. doi: 10.1016/j.ajhg.2019.02.008.

K. A. Lawrence, T. Gjorgjieva, D. Nachun, and S. B. Montgomery. Focus on single-gene effects limits discovery and interpretation of complex-trait-associated variants. *The American Journal of Human Genetics*, 113(4):842–851, 2026. doi: 10.1016/j.ajhg.2026.02.022.

X. Liu, X. Chen, B. Zhong, et al. Transcription factor achaete-scute homologue 2 initiates follicular T-helper-cell development. *Nature*, 507:513–518, 2014. doi: 10.1038/nature12910.

Y. Liu, S. Chen, Z. Li, A. C. Morrison, E. Boerwinkle, and X. Lin. ACAT: A fast and powerful p value combination method for rare-variant analysis in sequencing studies. *The American Journal of Human Genetics*, 104(3):410–421, 2019. doi: 10.1016/j.ajhg.2019.01.002.

S. D. Lombardo, E. Mazzon, M. S. Basile, G. Campo, F. Nicoletti, and P. Fagone. Modulation of Tetraspanin 32 (TSPAN32) expression in T cell-mediated immune responses and in multiple sclerosis. *International Journal of Molecular Sciences*, 20(18):4323, 2019. doi: 10.3390/ijms2018 4323.

L. R. O'Sullivan, M. R. Cahill, and P. W. Young. The importance of alpha-actinin proteins in platelet formation and function, and their causative role in congenital macrothrombocytopenia. *International Journal of Molecular Sciences*, 22(17):9363, 2021. doi: 10.3390/ijms22179363.

A. Saha and A. Battle. False positives in trans-eQTL and co-expression analyses arising from RNA-sequencing alignment errors. *F1000Research*, 7:1860, 2018. doi: 10.12688/f1000research.17145.2.

The 1000 Genomes Project Consortium. A global reference for human genetic variation. *Nature*, 526(7571):68–74, 2015. doi: 10.1038/nature15393.

W. Zhou et al. Efficient and accurate mixed model association tool for single-cell eQTL analysis. *medRxiv*, 2024. doi: 10.1101/2024.05.15.24307317. URL https://www.medrxiv.org/content/10.1101/2024.05.15.24307317v1. Preprint.

P. Zjablovskaja, M. Kardosova, P. Danek, et al. EVI2B is a C/EBP alpha target gene required for granulocytic differentiation and functionality of hematopoietic progenitors. *Cell Death and Differentiation*, 24(4):705–716, 2017. doi: 10.1038/cdd.2017.6.
